# Comparative genomic and transcriptomic analyses of Strongylus vulgaris reveal developmental and evolutionary deployment of parasitism in a migratory equine nematode

**DOI:** 10.64898/2026.07.17.739161

**Authors:** Nichol E. Ripley, Britt Ripley, Kai Li, Elizabeth Hudson, Daniel K. Howe, Theodore Kalbfleisch, Melissa L. Smith, Martin K. Nielsen

## Abstract

**Background:** *Strongylus vulgaris* is a highly pathogenic equine strongyle whose larval stages migrate through the mesenteric arterial system, yet the molecular basis of its development, host association, and evolutionary biology remains poorly resolved. We generated an integrated genomic and transcriptomic resource to characterize genome structure, gene annotation, stage-associated expression, candidate secreted proteins, isoform diversity, gene-family evolution, putative horizontal gene transfer, and drug resistance-associated homologs in *S. vulgaris*.

**Results:** PacBio HiFi sequencing produced a 329.0 Mb genome assembly comprising 3,913 contigs, with a contig N50 of 147 kb and 94.8% BUSCO completeness, substantially improving the prior fragmented draft. Repeat annotation identified 127.5 Mb of repetitive sequence, representing 38.72% of the genome and dominated by unclassified repeats. Integration of RNA-seq-guided annotation with PacBio Iso-Seq evidence refined 16,387 loci and 24,546 transcripts, generating an isoform-retaining discovery proteome of 23,363 predicted proteins. Functional annotation supported 93.5% of predicted proteins and identified 1,530 unknown or weakly annotated candidates. Consensus secretome prediction identified 2,210 high-confidence putative secreted proteins. Stage-associated transcriptomics showed that development was the dominant axis of expression variation, with 5,299 genes differentially expressed between larvae and adults and strong adult sex-associated divergence. Isoform analysis identified 490 high-confidence isoform switches, concentrated primarily in the ML5 female-to-adult female transition. Comparative genomics identified 27,641 orthogroups, 201 *S. vulgaris*-specific orthogroups, and, after repeat-aware filtering, 48 expanded and 250 contracted gene families. Structure-guided annotation prioritized migratory-stage-enriched secreted candidates, including Cysteine-rich secretory proteins, Antigen 5, and Pathogenesis-related 1 (CAP), Sperm-coating protein (SCP) Tpx-1/Ag5/PR-1/Sc7 protein superfamily (TAPS) -like, von Willebrand factor type A (VWA) - domain, lipid-binding-like, DNase II-like, and peptidase-like proteins. Conservative screening retained 17 putative horizontal gene transfer (HGT) candidates, and drug resistance-homolog analysis recovered 18 nonredundant *S. vulgaris* homologs without canonical β-tubulin benzimidazole-resistance substitutions.

**Conclusions:** These results establish the first integrated, high-quality molecular framework for *S. vulgaris* and show that its parasitic biology is developmentally structured, isoform-rich, and shaped by both conserved strongylid features and lineage-specific gene-family change. This resource provides a foundation for future studies of interhost migration, host interaction, parasite evolution, and genomic surveillance.

## Background

Gastrointestinal nematodes remain a major threat to equine health and parasite-control programs, with cyathostomins and large strongyles differing substantially in life history, tissue tropism, and pathogenic potential [1–3]. Among the large strongyles, *S. vulgaris* is of particular concern because its larval stages undergo extraintestinal migration through the cranial mesenteric arterial system before returning to the large intestine as adults [4–7]. This arterial migratory phase is central to the parasite’s clinical significance and distinguishes *S. vulgaris* from cyathostomins, whose pathology is primarily associated with larval development in the intestinal wall [3,4,7]. Although intensive anthelmintic treatment historically reduced *S. vulgaris* prevalence, modern surveillance-based and selective-treatment programs have renewed concern that this highly pathogenic species may persist or re-emerge when species-specific monitoring is insufficient [1,7,8].

Despite its clinical and biological importance, the molecular biology of *S. vulgaris* remains poorly resolved. The previously available *S. vulgaris* genome was generated from short-read data and was highly fragmented, limiting analyses that depend on accurate gene structure, repeat resolution, transcript-supported annotation, and comparative gene-family inference [9]. In other parasitic nematodes, improved genome and transcriptome resources have provided essential frameworks for investigating development, host interaction, genome evolution, and drug-response biology [9–13]. A more complete and transcript-supported genomic resource is therefore needed to determine how genome architecture, coding-gene content, and regulatory complexity contribute to the biology of this migratory equine strongyle. The life cycle of *S. vulgaris* involves major transitions among tissue-associated larval migration, intravascular development, intestinal adult maturation, and reproduction [4,7]. These transitions raise the question of whether the parasite deploys a generalized parasitic program or distinct molecular repertoires across developmental stages and sexes. Short-read RNA-seq can identify stage- and sex-associated expression patterns, whereas long-read transcript sequencing improves gene-model refinement and resolution of transcript isoforms [14,15]. Isoform diversity may be especially important in non-model parasites because alternative transcript usage can alter coding sequence, transcript structure, and regulatory potential [16,17]. In parallel, predicted secreted proteins provide a focused candidate set for host-interface biology, because helminth secretomes include extracellular molecules involved in invasion, migration, feeding, immune exposure, and host interaction [18–20].

Placing these molecular features in an evolutionary context is also necessary for distinguishing conserved strongylid biology from lineage-specific features of *S. vulgaris*. Comparative genomics can identify shared orthology, species-restricted genes, and gene-family expansions or contractions that may reflect developmental, ecological, or host-associated divergence [9,10,13]. In addition, exploratory screening for putative horizontal gene transfer can identify unusual loci with non-metazoan similarity patterns, although such candidates require conservative interpretation and additional validation [21–23]. Finally, an improved genome provides a baseline for identifying homologs of resistance-associated genes characterized in other nematodes. Because homolog detection is not equivalent to phenotypic resistance, such loci are best interpreted as a reference framework for future surveillance rather than as evidence of resistance in *S. vulgaris* [24–27].

Here, we present an integrated genomic and transcriptomic analysis of *S. vulgaris* using PacBio HiFi genome sequencing, PacBio Iso-Seq transcript sequencing, and Illumina RNA-seq across major parasitic stages. We use these resources to generate an improved reference assembly, refine gene and transcript models, characterize repeat content and protein annotation, identify stage-associated expression and isoform-switching patterns, predict candidate secreted proteins, evaluate comparative orthology and gene-family evolution, screen conservatively for putative horizontal gene transfer candidates, and establish a baseline catalog of resistance-associated homologs. Together, these analyses provide a molecular framework for investigating development, host association, evolutionary divergence, and future genomic surveillance in this pathogenic migratory equine nematode.

## Methods

### Parasite recovery, life-stage identification, and ethical approval

All *S. vulgaris* specimens used in this study were recovered during necropsy from 11 humanely euthanized foals, 5–9 months of age, from the University of Kentucky parasitology research herd. This closed herd has not received anthelmintic treatment since 1979. Foals were of mixed Thoroughbred lineage and were born in 2021 or 2022. Euthanasia and necropsy were conducted under approval from the University of Kentucky Institutional Animal Care and Use Committee (protocol #2021-3879) and in accordance with accepted veterinary euthanasia guidelines [28].

Parasites were collected from the cranial mesenteric arteries (CMAs), intestinal lumen, mucosal surface, and intestinal wall cysts. Fourth-stage larvae (L4) were collected from the CMA. Migratory fifth-stage larvae (ML5) were collected from the CMA and from developing cysts in the intestinal wall. adult worms were collected from the intestinal lumen and mucosal surface of the large intestine. Following recovery, parasites were maintained in phosphate-buffered saline until processing.

Life stage and species identity were assigned morphologically using established taxonomic keys [29]. Adult worms and ML5 larvae were sexed where morphological features permitted.

### Nucleic acid extraction, quality control, library preparation, and sequencing

Genomic DNA was extracted from one individual ML5 female larva and from a pool of three ML5 female larvae using the MagAttract High Molecular Weight gDNA Kit (Qiagen, Valencia, CA, USA), with final elution in 50 µL nuclease-free water. The individual ML5 female DNA extraction was used for the primary genome assembly library, and the pooled ML5 female DNA extraction was used to generate an additional long-read dataset for assembly refinement.

Total RNA was extracted using the RNeasy Mini Kit (Qiagen), with final elution in 50 µL nuclease-free water. Preliminary RNA extractions were used to determine the number of individuals required to obtain sufficient RNA from each life stage. For long-read transcript sequencing, RNA was extracted from one pool of 45 L4 larvae and one pool of 10 ML5 female larvae. For short-read RNA-seq, RNA was extracted from replicate samples representing L4, ML5, and adult stages. L4 samples contained 45 larvae per replicate from mixed foals, whereas ML5 and adult samples contained 10 individuals per replicate from mixed foals and were subdivided by sex and collection site where applicable. After quality control, 13 RNA samples were advanced to Illumina library preparation.

Nucleic acid concentration and purity were assessed using Qubit fluorometry (Invitrogen/Thermo Fisher Scientific, Waltham, MA, USA) and NanoDrop spectrophotometry (Thermo Fisher Scientific). DNA fragment size and RNA integrity were assessed using Bioanalyzer instruments (Agilent Technologies, Santa Clara, CA, USA). Samples were advanced to library preparation only when they met platform-specific input and quality requirements.

Two PacBio HiFi genomic libraries were prepared by the Sequencing Technology Center at the University of Louisville (Louisville, KY, USA). The primary genome assembly library was generated from 18 ng of genomic DNA from a single ML5 female using the PacBio SMRTbell HiFi ultra-low DNA input workflow. A second PacBio HiFi whole-genome library was prepared from pooled ML5 female DNA using the SMRTbell HiFi WGS Prep Kit 1.0. Libraries were size-selected and quality checked prior to sequencing.

For full-length transcript sequencing, two PacBio Iso-Seq libraries were prepared from pooled L4 and ML5 female RNA using the Iso-Seq v2 workflow with SMRTbell prep kit 3.0 and 300 ng total RNA input. Iso-Seq libraries were sequenced on the PacBio Sequel IIe platform.

For short-read transcriptome profiling, 13 stranded total RNA libraries were prepared using the Illumina Stranded Total RNA Prep with Ribo-Zero Plus chemistry and sequenced on an Illumina NextSeq 2000 platform using a P2 flow cell with 1 × 100 bp reads. Because parasite material was collected from host tissues, custom ribodepletion probes targeting both *S. vulgaris* and equine ribosomal transcripts were incorporated during library preparation. Together, the 13 Illumina RNA-seq libraries yielded 57 Gb of raw sequence data.

### Genome size estimation, host decontamination, assembly, QC, and repeat annotation

Genome assembly was performed using PacBio HiFi reads generated from the single ML5 female genomic DNA library. A second PacBio HiFi dataset generated from pooled ML5 female larvae was retained for assembly refinement and read-support assessment rather than for primary assembly construction.

Raw HiFi reads from the single-worm dataset were first screened for residual PacBio adapter sequence using HiFiAdapterFilt [30]. To reduce host contamination, reads were aligned to the *Equus caballus* TB-T2T reference genome [31] using minimap2, and unmapped reads were retained as the parasite-enriched assembly input. The retained read set was then screened with Kraken2 to identify and remove residual non-target contamination prior to assembly.

Genome size was estimated from 21-mer frequencies generated from the parasite-enriched single-ML5-female HiFi dataset using Merqury and size estimation tool GenomeScope2 [32,33]. The host-depleted and contaminant-filtered reads were assembled *de novo* using Hifiasm [34]. The resulting assembly was evaluated for total span, contig number, contig N50, longest contig, GC content, and read support.

Genome polishing and coverage assessment were performed using the pooled ML5 female HiFi dataset. Pooled reads were aligned back to the selected assembly with minimap2 to evaluate read support and coverage consistency. Residual contamination and assembly composition were further assessed using BlobTools2, incorporating GC content, read coverage, and taxonomic assignment.

Assembly completeness was evaluated using BUSCO v5.5 [35] with the nematoda_odb10 lineage dataset. The final cleaned assembly was carried forward for repeat annotation, masking, and evidence-guided gene prediction.

Repetitive elements were annotated using a species-specific repeat-discovery workflow. RepeatModeler2 was used to generate a *de novo* repeat library, and RepeatMasker [36] was then used to classify and soft-mask repetitive sequences in the assembly. The resulting soft-masked genome was used for downstream structural annotation, and major repeat classes were summarized to characterize genome repeat composition.

### Alignment of short-read RNA-seq and long-read Iso-Seq data, structural annotation, and transcript model construction

Transcript-supported structural annotation was performed using both short-read Illumina RNA-seq and long-read PacBio Iso-Seq evidence generated from major parasitic stages of *S. vulgaris*. The Illumina RNA-seq dataset included 13 stranded total RNA libraries representing L4, ML5 female, ML5 male, adult female, and adult male samples. The Iso-Seq dataset included two full-length transcript libraries generated from pooled L4 and pooled ML5 female material.

Short-read RNA-seq reads were aligned to the soft-masked *S. vulgaris* genome using STAR, and the resulting alignments were used as transcript evidence for genome annotation. Structural gene prediction was performed with BRAKER3 in ETP mode using RNA-seq evidence and a curated OrthoDB Nematoda protein database as external protein hints [37]. The protein-hint database contained 277,101 sequences and included *Caenorhabditis elegans* proteins to support prediction of conserved nematode loci. Within BRAKER3, GeneMark-ETP generated evidence-supported gene predictions, AUGUSTUS performed trained gene prediction, and TSEBRA consolidated the resulting models into a final annotation set.

Iso-Seq reads were aligned to the soft-masked genome using minimap2 with parameters “-ax splice:hq -uf”. The primary alignment bam file was used in subsequent transcript model integration and annotation refinement analyses. BRAKER3 gene models and Iso-Seq-supported transcript models were merged using StringTie v2 in merge mode, with BRAKER3 models retained as the primary coordinate framework. Iso-Seq evidence was used to support splice-junction validation, refine exon-intron structures, and extend transcript boundaries where supported by full-length transcript data.

The merged transcript annotation was converted to GFF3 format using GFFRead for downstream compatibility. Structural concordance between the original BRAKER3 annotation and the refined transcript set was assessed using GFFCompare. A gene-to-transcript crosswalk table was generated from the final merged annotation to support downstream integration of gene, transcript, and protein identifiers.

Completeness of the refined transcript-supported annotation was evaluated using BUSCO with the nematoda_odb10 lineage dataset. Because the final annotation retained isoform diversity, duplicated BUSCO calls were interpreted in the context of transcript-level redundancy rather than as direct evidence of assembly duplication.

### Proteome construction, representative proteome filtering, and functional annotation

Protein prediction was performed from the refined transcript-supported annotation. Open reading frames were identified using TransDecoder, and predicted proteins were evaluated using homology- and domain-based evidence. Candidate protein sequences were searched against UniProt Swiss-Prot using BLASTP or DIAMOND, and conserved domains were identified using HMMER hmmscan against Pfam-A. Predicted protein identifiers were linked back to transcript and gene identifiers using a gene–transcript–protein crosswalk to preserve coordinate and annotation provenance.

Three proteome datasets were generated for downstream analyses. First, a discovery proteome was created from all valid TransDecoder-supported protein isoforms. This isoform-retaining proteome was used for functional annotation, secretome prediction, structure-guided analysis of weakly annotated proteins, and other candidate-gene analyses where transcript diversity could be biologically informative. Second, a representative proteome was generated by collapsing the discovery proteome to a single protein per parent locus, retaining the longest predicted protein isoform. This nonredundant proteome was used for gene-level annotation summaries and to reduce inflation from multiple isoforms of the same locus. Third, a comparative proteome was generated from the representative proteome for orthology and gene-family analyses. Proteins with repeat- or transposable element-associated domain signatures were removed before OrthoFinder and CAFE5 analyses to reduce inflation of gene-family estimates by repeat-derived proteins [38,39]. The final comparative proteome was therefore used only for orthology inference and gene-family evolution analyses, whereas the broader discovery proteome was retained for functional and candidate-protein analyses.

Functional annotation was performed using an integrated homology-, domain-, and pathway-based workflow. Protein sequences were searched against UniProt Swiss-Prot and a nematode-filtered UniProtKB/TrEMBL database using DIAMOND. The nematode TrEMBL database was used as a lineage-focused homology resource to complement curated Swiss-Prot annotations. Pfam domain annotations were incorporated from hmmscan results, and KEGG Orthology and pathway assignments were generated using KAAS [40]. Swiss-Prot, TrEMBL, Pfam, and KEGG outputs were merged into a master annotation table linking gene, transcript, and protein identifiers with available homology, domain, and pathway evidence. Proteins lacking strong homology or conserved-domain support were retained as an unknown or weakly annotated candidate set for downstream prioritization, including secretome-linked, expression-linked, and structure-guided analyses.

### Candidate secretome prediction

Candidate secreted proteins were predicted from the discovery proteome to retain biologically plausible isoforms that could be relevant to host-interaction analyses. A consensus filtering workflow was used to identify proteins with features consistent with soluble extracellular secretion. SignalP 6.0 and TargetP 2.0 were used to identify proteins with N-terminal secretion signals and to exclude likely mitochondrial-targeted proteins [41]. Phobius 1.01 and TMHMM 2.0 were used to evaluate membrane topology and remove proteins with internal transmembrane domains inconsistent with soluble secretion [42].

Predictor outputs were integrated to generate a high-confidence candidate secretome. Proteins retained in the final secretome set were required to have secretion-signal support and no conflicting topology evidence suggestive of membrane anchoring. The resulting candidate secretome was cross-referenced with differential expression results, particularly the larvae vs adult contrast, to identify secreted proteins with stage-associated expression patterns.

### Differential expression, enrichment, and pathway analysis

Differential gene expression analyses were performed using the 13 retained Illumina RNA-seq libraries representing L4, ML5 female, ML5 male, adult female, and adult male samples. Gene-level counts were generated from RNA-seq alignments using featureCounts from the subread package, with the final transcript-supported genome annotation used as the counting reference.

Raw count matrices were analyzed in DESeq2 using the standard negative binomial framework. Size-factor normalization was applied prior to differential expression testing. Pairwise contrasts were performed among biologically relevant sample groups: adult male vs adult female, larvae vs adult, ML5 female vs adult female, ML5 male vs adult male, and ML5 male vs ML5 female. Genes with Benjamini–Hochberg adjusted *P* values ≤ 0.05 were considered significantly differentially expressed [43]. Log2 fold-change shrinkage was not applied, so reported fold changes represent unshrunken DESeq2 estimates.

To support biological interpretation, differential expression tables were merged with functional annotations from the master annotation table. Gene names and functional descriptions were assigned using a prioritized annotation framework incorporating Swiss-Prot, nematode homology, Pfam/domain evidence, and KEGG assignments where available.

Global expression patterns were evaluated using principal component analysis and hierarchical clustering of the most variable genes. Differential expression results were visualized using volcano plots and MA plots for each contrast. Functional enrichment analyses were performed in R using clusterProfiler, with Gene Ontology and KEGG-linked annotations derived from the integrated annotation workflow [44]. Enrichment analyses were performed on contrast-specific DEG sets, and significance was assessed using Benjamini–Hochberg multiple-testing correction. KEGG pathway results were interpreted conservatively. Pathway labels derived from vertebrate disease or immune-system terminology were not treated as direct evidence of equivalent parasite biology but instead were evaluated through inspection of the underlying driver genes and interpreted as conserved molecular modules where appropriate.

### Alternative splicing and isoform-switching analyses

Alternative splicing and differential isoform usage were evaluated using transcript models from the refined annotation. Isoform-level abundance and count matrices were generated from StringTie-derived transcript estimates, and transcript-to-gene relationships were defined using the final gene–transcript mapping table.

Differential isoform usage was analyzed in R using IsoformSwitchAnalyzeR with DEXSeq as the statistical framework [45]. Because replicated groups are required for isoform-switch testing, the unreplicated L4 group was excluded from differential isoform usage analyses. Low- and zero-expression transcripts were filtered prior to testing to reduce the multiple-testing burden. Pairwise isoform-usage comparisons were then performed among replicated groups, including adult male vs adult female, ML5 female vs adult female, ML5 male vs adult male, and ML5 male vs ML5 female.

Open reading frames were predicted from the refined transcript sequences to support interpretation of coding potential and transcript consequences. Alternative splicing events were classified within IsoformSwitchAnalyzeR, including exon skipping, intron retention, mutually exclusive exons, alternative 5′ and 3′ splice-site usage, and alternative transcription start and termination site usage. Representative isoform-switching loci were visualized using transcript-structure plots, with emphasis on switches affecting coding structure or predicted nonsense-mediated decay sensitivity.

### Orthology inference, phylogenomics, and gene-family evolution

Orthology inference was performed using the filtered comparative proteome to minimize inflation from alternative isoforms and repeat-associated proteins. Protein sets from *S. vulgaris* and comparison clade V nematode species (*C. elegans, Nippostrongylus brasiliensis, Haemonchus contortus, Oesophagostomum dentatum, Cylicocyclus nassatus, Necator americanus,* and *Ancylostoma ceylanicum*) were analyzed with OrthoFinder to identify orthogroups, infer single-copy orthologues, and generate the species phylogeny used for downstream comparative analyses.

Gene-family expansion and contraction were modeled using CAFE5 with the OrthoFinder orthogroup count matrix and rooted species tree as inputs. To account for potential assembly or annotation error, the CAFE5 error model was used to estimate a global error distribution. A single-lambda model was then fitted to estimate the genome-wide birth–death rate of gene gain and loss across the sampled nematode phylogeny.

To reduce artifacts from repetitive elements and extreme copy-number inflation, orthogroups were filtered prior to biological interpretation. Families dominated by transposable element- or repeat-associated Pfam domains, including transposase, retrotransposon, gag, or polymerase-like domains, were excluded from interpreted expansion results. Gene families were considered significantly expanded or contracted when they showed family-wide *P* ≤ 0.05 in CAFE5.

For prioritized expanded families, functional interpretation was based on consensus evidence across member proteins, including Swiss-Prot similarity, nematode homology, Pfam domains, and KEGG annotations where available. Families lacking consistent functional support were retained as lineage-specific or weakly annotated candidates rather than assigned speculative functions.

### Structure-guided functional annotation of prioritized candidate proteins

Structure-guided annotation was used to improve interpretation of proteins that remained unknown or weakly annotated after conventional homology-, domain-, and pathway-based annotation. Candidate proteins were selected from the discovery proteome to retain isoforms that could be biologically informative. Proteins were prioritized for structural analysis if they met one or more criteria, including predicted secretion, migratory-stage differential expression, membership in expanded or lineage-specific gene families, or limited conventional annotation despite transcript or protein-level support.

Generated AlphaFold2 models were evaluated for structural interpretability using model-confidence metrics, including mean predicted local distance difference test scores, per-residue confidence, predicted aligned error, and domain architecture. Proteins were classified as high-confidence, partial-confidence, or low-confidence structural models. For multi-domain proteins with confident domains but uncertain inter-domain orientation, domains were interpreted separately where appropriate.

Structural similarity searches were performed using Foldseek against experimentally resolved structures in the Protein Data Bank and, where informative, AlphaFoldDB [46–48]. Structural matches were interpreted conservatively, with greatest weight given to high-coverage matches to proteins with coherent functional annotations. Matches limited to short regions, broad structural superfamilies, or functionally inconsistent hits were treated as fold-level evidence only.

DeepFRI was used as a complementary structure-based functional prediction tool. Predicted Gene Ontology terms and confidence scores were evaluated alongside Foldseek results, model confidence, domain evidence, predicted secretion status, and expression pattern. DeepFRI predictions were used as supporting evidence rather than as standalone functional assignments.

For selected high-priority proteins with structural novelty, limited sequence-based annotation, or potentially divergent Foldseek results, DALI was used as an additional structural comparison method. DALI-supported interpretations were retained only when they were consistent with model confidence, structural coverage, and broader biological evidence.

### Horizontal gene transfer candidate assessment

Putative horizontal gene transfer (HGT) candidates were evaluated using a conservative multi-step workflow designed to distinguish candidate foreign-derived loci from contamination, annotation artifacts, and deeply conserved genes with uneven database representation. Analyses were performed using the predicted *S. vulgaris* proteome, final genome annotation, RNA-seq expression data, and genome assembly.

All predicted *S. vulgaris* proteins were searched against a locally formatted, taxonomy-aware NCBI nr protein database using DIAMOND blastp. For each query, the top 50 hits were retained using an E-value threshold of < 1 × 10^-5^. Hits were parsed with associated taxonomic identifiers and assigned to higher-level evolutionary lineages. Alien Index scoring was then used to identify proteins with stronger similarity to non-metazoan sequences than to metazoan sequences. Because ancient HGT events may be shared among nematodes, Nematoda was masked during Alien Index calculation to avoid suppressing candidates with nematode homologs that may themselves reflect older acquisition events.

Candidate proteins were subjected to strict filtering to reduce contamination-style false positives. Retained candidates were required to have an Alien Index ≥ 45, mean RNA-seq expression support ≥ 5 counts, placement on scaffolds longer than 1 kb, and < 70% amino acid identity to the best non-metazoan match. Candidates failing these criteria were excluded from the strict HGT candidate set.

For retained candidates, genomic context was evaluated using scaffold identity, scaffold length, gene coordinates, neighboring gene annotations where available, transcript support, and coding-sequence structure. Expression support was assessed from RNA-seq count data, and candidates lacking expression evidence were excluded. These filters were used to reduce the likelihood that retained candidates represented unexpressed contaminants, short scaffold artifacts, or unsupported gene models.

Phylogenetic validation was performed for the strict candidate set. Homologous sequences were retrieved for each candidate, including top non-metazoan matches and relevant nematode or metazoan homologs where available. Protein sequences were aligned using MAFFT, and maximum-likelihood phylogenies were inferred using IQ-TREE2 with 1,000 ultrafast bootstrap replicates [49]. Candidate phylogenies were interpreted conservatively, with stronger support assigned only when *S. vulgaris* candidates grouped more closely with non-metazoan sequences than expected under ordinary nematode inheritance.

Codon-composition analysis was used as an additional line of evidence. Coding sequences were extracted from the final genome annotation, and codon-frequency profiles were calculated for the full gene set and for the strict HGT candidates. Principal component analysis was then used to compare candidate codon-usage profiles against the broader native coding background. HGT candidates were interpreted conservatively and discussed as putative only when supported by multiple lines of evidence, including alienness score, expression support, gene structure, scaffold context, phylogenetic placement, and coding-sequence composition.

### Baseline survey of resistance-associated homologs

Resistance-associated homologs were surveyed in *S. vulgaris* as a baseline comparative analysis rather than as a test of phenotypic resistance. A curated bait set was generated from resistance-associated genes previously described in *C. elegans*, cyathostomins, and other strongylid or animal-parasitic nematodes. After removal of redundant entries, 43 unique resistance-associated protein queries were retained for screening.

To identify corresponding *S. vulgaris* homologs, curated bait sequences were searched against both the representative *S. vulgaris* proteome and the genome assembly. Protein-level homologs were identified using DIAMOND searches against the representative proteome, and genome-level matches were identified using tBLASTn searches against the genome assembly. Candidate homologs were retained using a threshold framework requiring E-value < 1 × 10^-10^, amino acid identity > 40%, and aligned sequence length > 150 amino acids.

Candidate homologs were mapped back to transcript and gene-structure information to recover genomic coordinates and exon metrics. Coordinate extraction was performed from the final genome annotation, generating BED-formatted loci and exon-structure summaries for downstream visualization and comparison.

To evaluate copy-number patterns and evolutionary relationships, full-length amino acid sequences were extracted for proteins assigned to targeted resistance-associated orthogroups across the comparative nematode dataset. Each orthogroup was aligned independently using MAFFT, and maximum-likelihood gene trees were generated using FastTree. Orthogroup-specific trees were used to assess paralogy, relative copy number, sequence divergence, and placement of *S. vulgaris* homologs relative to homologs from comparison nematode species.

Because this analysis did not include phenotypically resistant isolates, treatment-selected material, or experimental drug-response validation, identified loci were interpreted as resistance-associated homologs rather than resistance markers. The resulting dataset provides a genomic baseline for future population-level surveillance, structural comparison, and targeted validation of candidate drug-response loci in *S. vulgaris*. A full flowchart of analyses is available in Additional file 1.

### Use of large language models

Large language models were used to assist with additional file data organization (ChatGPT Pro v5.5; Open AI, San Francisco, CA, USA) and bioinformatic and statistical coding support (Gemini Pro v3.1; Google, New York, NY, USA). All scientific analyses, data interpretation, conclusions, and final manuscript text were independently reviewed, verified, and approved by the authors.

## Results

### Sequencing and read filtering generated curated genome and transcriptome datasets across major parasitic stages

Genome and transcriptome datasets were generated from *S. vulgaris* life stages selected to support genome assembly, gene-model refinement, and developmental expression analyses. Genome assembly used PacBio HiFi reads from a single ML5 female larva, with an additional pooled ML5 female HiFi dataset generated for assembly refinement and read-depth assessment. After adapter, host, and contaminant filtering, the single-worm dataset retained approximately 2.9 million parasite-enriched HiFi reads, providing an estimated 63.3× assembly depth.

Transcriptome resources included PacBio Iso-Seq libraries from pooled L4 and ML5 female material, yielding approximately 131,000 and 160,000 full-length non-concatemer reads, respectively. Short-read RNA-seq was generated from 13 retained Illumina libraries spanning L4, ML5 female, ML5 male, adult female, and adult male samples. Individual RNA-seq libraries contained 35.4–48.3 million reads, with a mean of 41.1 million reads per library. Together, these datasets provided long-read genomic coverage for assembly, full-length transcript evidence for annotation refinement, and stage- and sex-resolved RNA-seq data for differential expression and isoform analyses (Table 1).

**Table 1.**
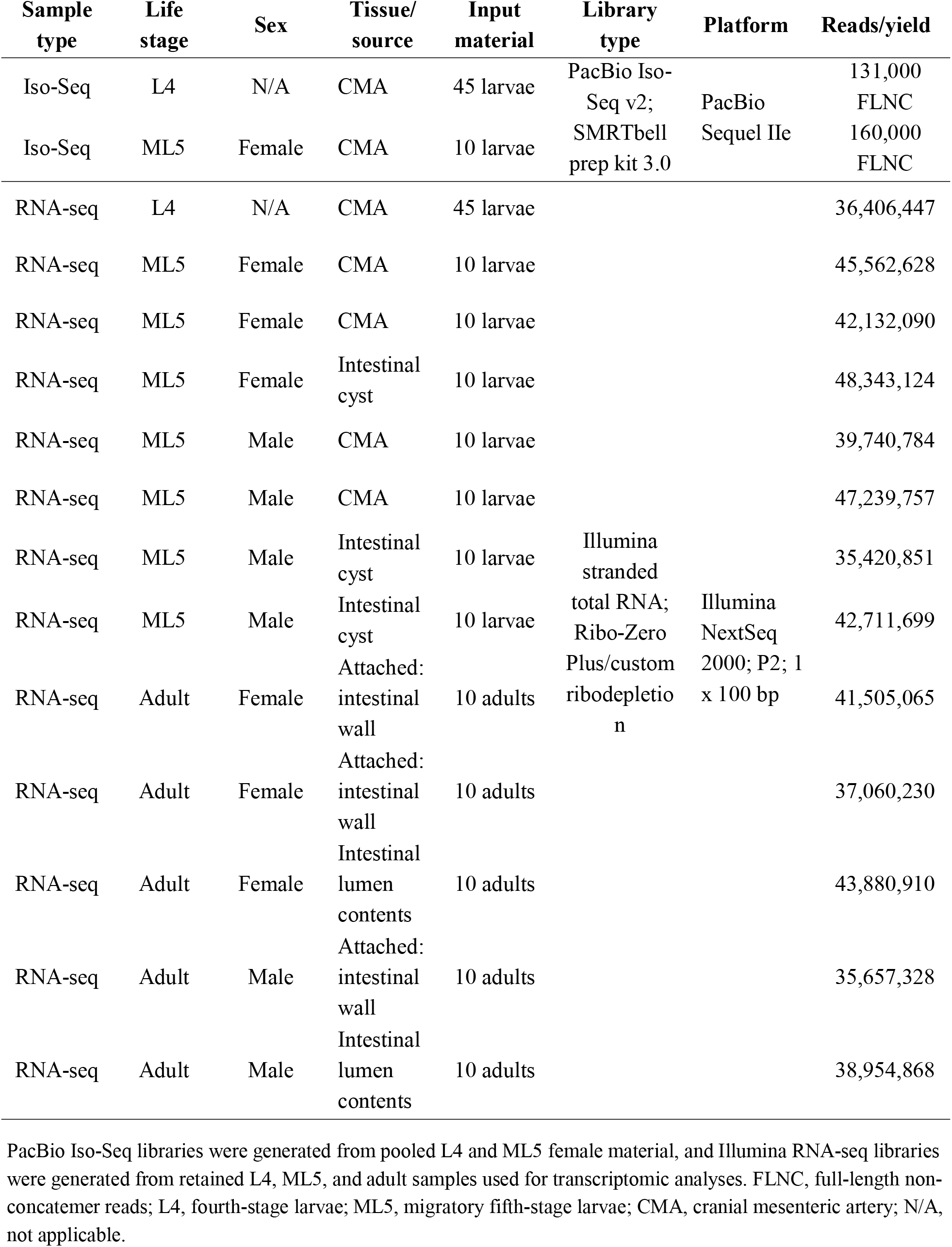
Sample and sequencing summary for transcriptomic datasets.

### PacBio HiFi assembly and refinement produced a substantially improved *S. vulgaris* **genome**

PacBio HiFi assembly of the parasite-enriched single-ML5-female dataset produced a final *S. vulgaris* genome assembly of 329.0 Mb. The assembly comprised 4,852 contigs, with a contig N50 of 142 kb, a longest contig of approximately 1 Mb, a mean contig length of 68 kb, and GC content of 38.7%.

BUSCO analysis against the nematoda_odb10 dataset recovered 94.8% complete BUSCOs, including 86.3% single-copy and 8.6% duplicated orthologs, indicating high assembly completeness and a substantial improvement over the previously published short-read draft genome. Assembly metrics and BUSCO comparisons are summarized in Figures 1A and 1B, with a full side-by-side assembly comparison provided in Additional file 2.

**Figure 1.**
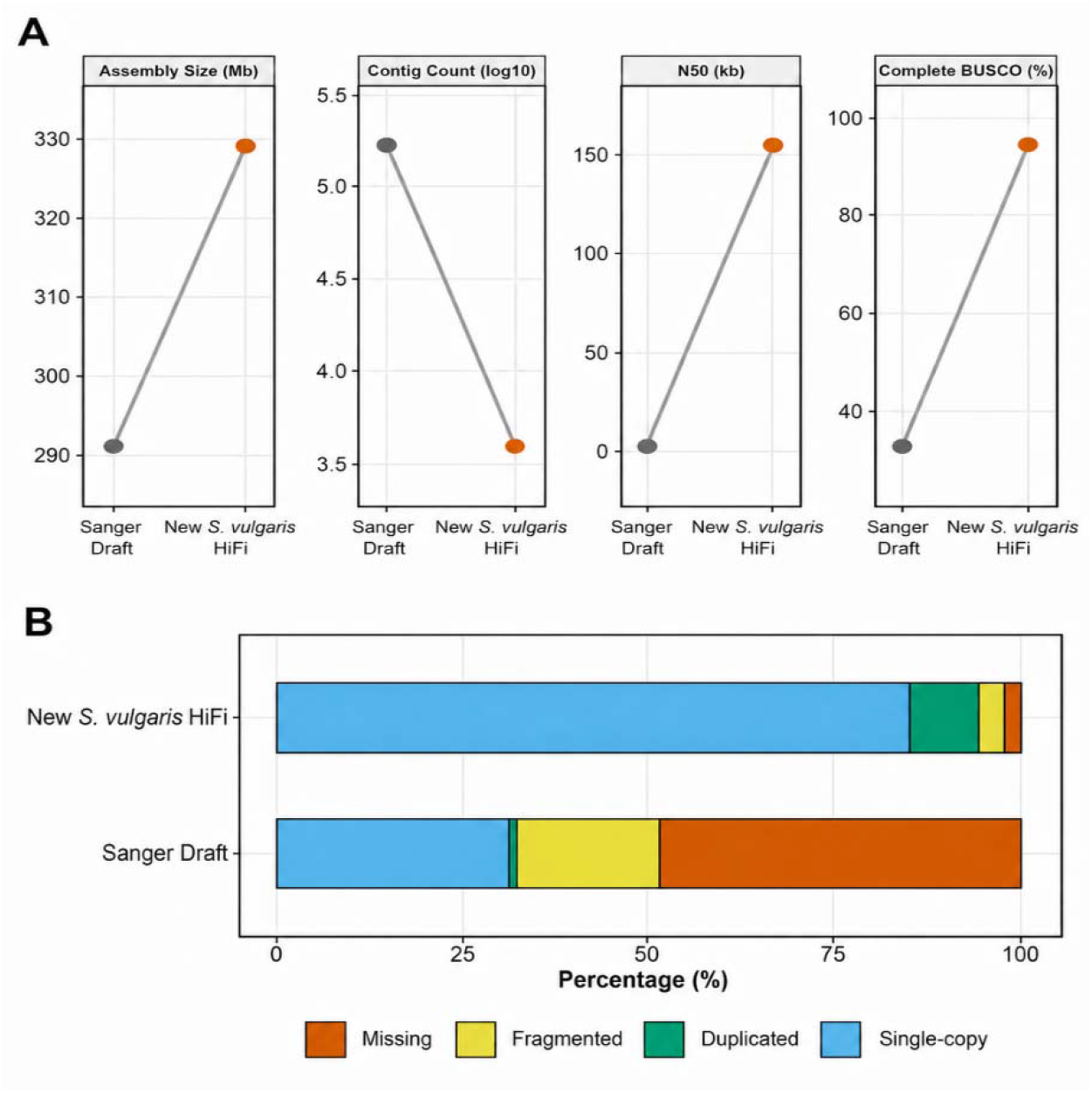
PacBio HiFi sequencing improved Strongylus vulgaris genome assembly contiguity and completeness. Comparison of the previously published S. vulgaris draft genome and the new PacBio HiFi assembly. (A) Assembly metrics showing genome size, contig count, contig N50, and complete BUSCO recovery. The HiFi assembly increased assembly size and contiguity while reducing contig number. (B) BUSCO completeness comparison using the nematoda_odb10 dataset. The HiFi assembly showed increased complete BUSCO recovery and reduced missing BUSCO content relative to the earlier draft assembly. BUSCO categories are shown as single-copy, duplicated, fragmented, and missing orthologs.

The pooled ML5 female HiFi dataset provided independent read support for the selected assembly. After host filtering, 6.8 million pooled reads aligned back to the assembly and supported an estimated read depth of 105.1×, without substantially altering assembly size or BUSCO completeness. Together, these results support the recovery of a substantially improved and highly complete reference assembly for S. vulgaris.

### Repeat annotation revealed a repeat-rich genome dominated by unclassified elements

Repeat annotation identified 127.5 Mb of repetitive sequence, corresponding to 38.72% of the S. vulgaris assembly. Most repeat content was assigned to unclassified elements, which accounted for more than one-quarter of the genome and represented the dominant repeat category (Figure 2). Among classified repeats, LINEs were the largest component, with smaller contributions from LTR elements, DNA transposons, simple repeats, small RNA-associated sequences, and low-complexity regions. Overall, the repeat landscape indicates that the S. vulgaris genome is repeat-rich and contains a large fraction of repetitive sequence that is either unclassified or highly diverged from currently characterized repeat families.

**Figure 2.**
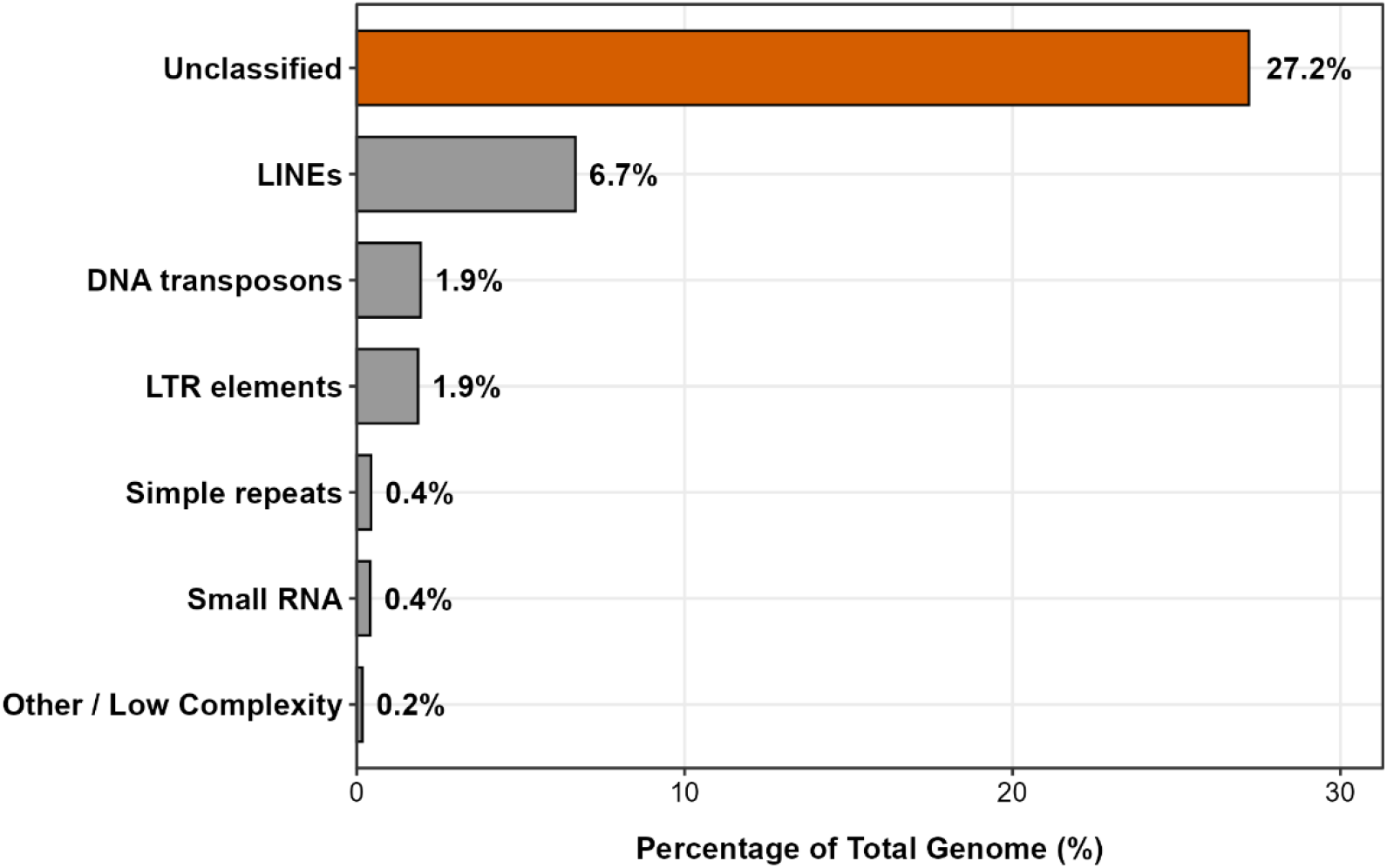
Repeat annotation revealed a repeat-rich S. vulgaris genome dominated by unclassified elements. Major repeat categories identified in the S. vulgaris genome assembly, shown as the percentage of total genom sequence.

### Transcript-supported annotation and proteome refinement generated curated gene and protein sets

Integration of RNA-seq-guided BRAKER3 annotation with long-read Iso-Seq evidence produced a refined S. vulgaris annotation containing 24,546 transcripts across 16,387 loci, including 4,607 multi-transcript loci. The refined annotation remained highly concordant with the original BRAKER3 gene models, with GFFCompare showing 99.1% transcript-level sensitivity, 99.8% locus-level sensitivity, 97.4% exon-level sensitivity, and 100.0% intron-level sensitivity.

Transcript-supported refinement also added structural information not captured in the initial annotation, including 1,259 novel loci and additional exon and intron structure. BUSCO analysi of the refined annotation recovered 90.9% complete nematode BUSCOs, with the elevated duplicated fraction consistent with retention of isoform diversity in the transcript-supported transcript set.

Protein prediction and filtering generated three coordinated proteome resources: an isoform-rich discovery proteome for candidate-protein analyses, a representative gene-level proteome, and a TE/repeat-filtered comparative proteome for orthology and gene-family analyses (Figure 3; Additional file 3). The final comparative proteome contained 16,385 proteins, providing the curated protein set used for downstream evolutionary comparisons.

**Figure 3.**
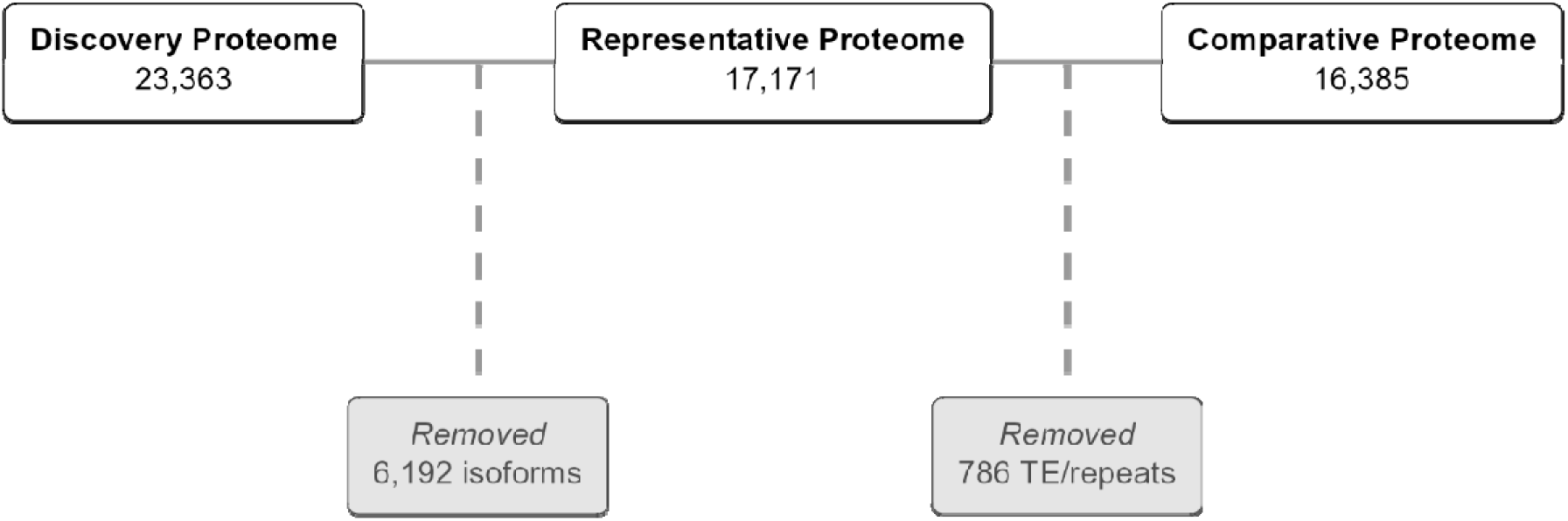
Proteome refinement generated coordinated discovery, representative, and comparative protein sets. Overview of the proteome refinement workflow used to generate protein resources for downstream analyses. The isoform-retaining discovery proteome contained 23,363 predicted proteins. Representative isoform selectio removed 6,192 isoform entries and produced a 17,171-protein representative proteome. Subsequent removal of repeat- and transposable element-associated proteins excluded 786 additional proteins and generated a final 16,385-protein comparative proteome for orthology and gene-family analyses.

### Functional annotation and KEGG pathway mapping characterized the predicted biological repertoire

Functional annotation of the S. vulgaris discovery proteome produced a master annotation table containing 23,363 protein entries. Integration of Swiss-Prot, nematode TrEMBL, Pfam, and KEGG evidence identified annotation support for most predicted proteins, with 21,833 entries receiving support from at least one source and 1,530 proteins retained as unknown or weakly annotated candidates (Figure 4).

**Figure 4.**
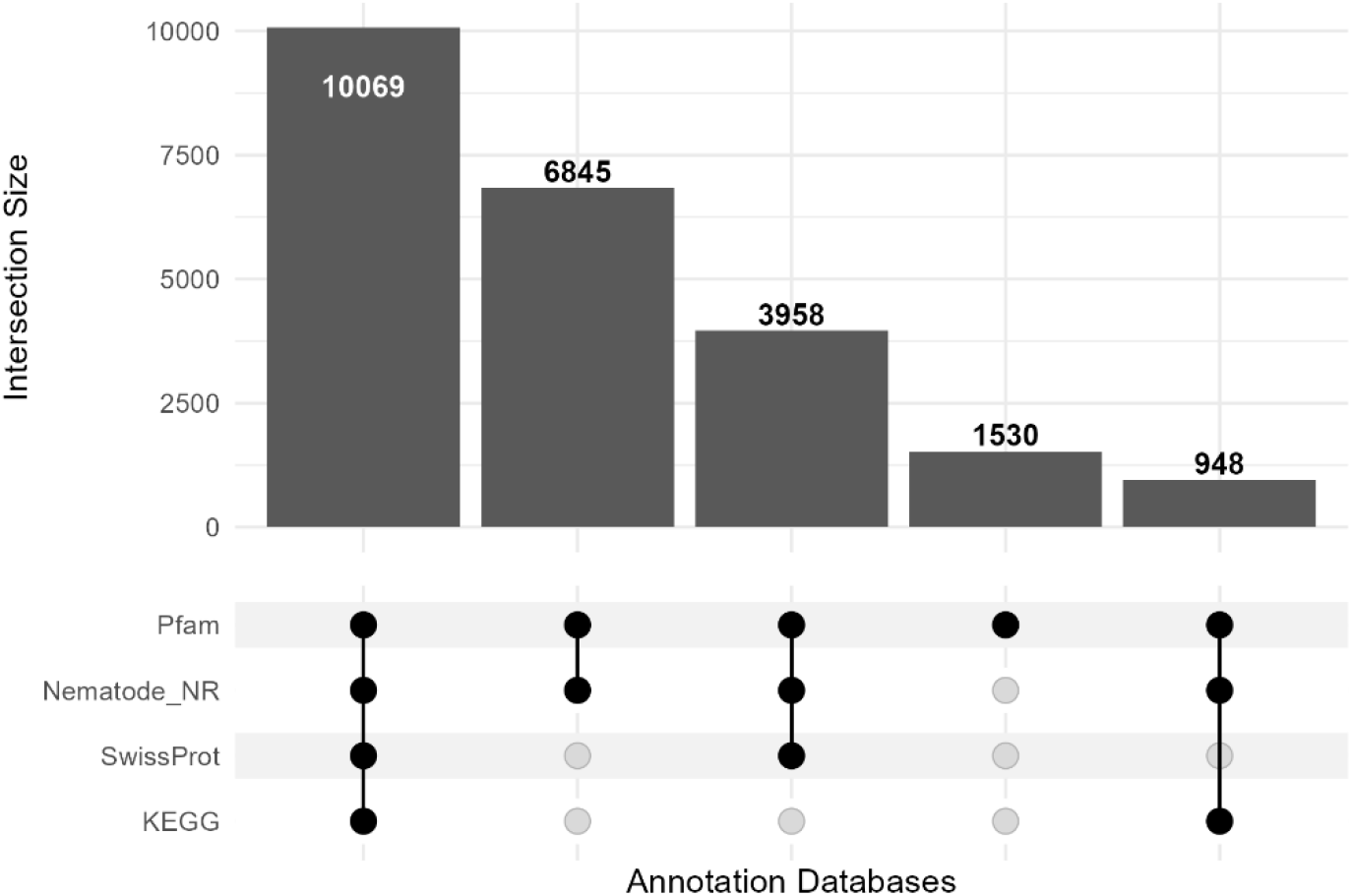
Integrated functional annotation characterized the predicted S. vulgaris proteome. UpSet-style summary of functional annotation support across Pfam, nematode nonredundant database searches, Swiss-Prot, and KEGG. Bars show the number of protein entries supported by each database intersection, and filled circles indicate the annotation sources contributing to each intersection.

At the representative gene level, 16,538 of 17,171 representative protein entries received putative functional support, corresponding to 96.3% of the representative annotation set. KEGG Orthology mapping assigned KO terms to 8,212 representative loci, and curated pathway mapping placed 4,963 loci into 428 KEGG pathways.

The most broadly represented pathway categories included core metabolism, lysosome-associated pathways, endocytosis, ribosome and spliceosome function, autophagy, cell adhesion, cytoskeletal regulation, oxidative phosphorylation, xenobiotic metabolism, and major signaling pathways. Together, these annotations support a gene repertoire consistent with developmental remodeling, cellular trafficking, environmental sensing, metabolic activity, stress tolerance, and host-interface biology.

Because KEGG pathway labels are derived largely from model-organism nomenclature, disease-or immune-labeled terms were interpreted conservatively as evidence of shared conserved gene modules rather than direct evidence of equivalent vertebrate disease or immune pathways in S. vulgaris.

### Secretome prediction identified candidate extracellular proteins with stage-associated expression

Consensus secretome prediction identified 2,210 high-confidence putative secreted proteins in the S. vulgaris discovery proteome, representing 9.5% of predicted proteins, see a full listing in Additional file 4. The remaining 21,153 proteins were classified as non-secreted after filtering for signal peptide support, subcellular targeting, and membrane topology. Thus, the predicted secretome represents a defined subset of the proteome with features consistent with extracellular deployment.

Integration of the candidate secretome with the Larvae vs Adult differential expression contrast showed that predicted secreted proteins were not uniformly expressed across development. The secretome included adult-biased proteins, larva-biased proteins, and proteins with more limited stage bias. Several labeled larva-biased secreted candidates, whereas the adult-biased set extended to proteins with larger positive fold changes (Figure 5A).

**Figure 5.**
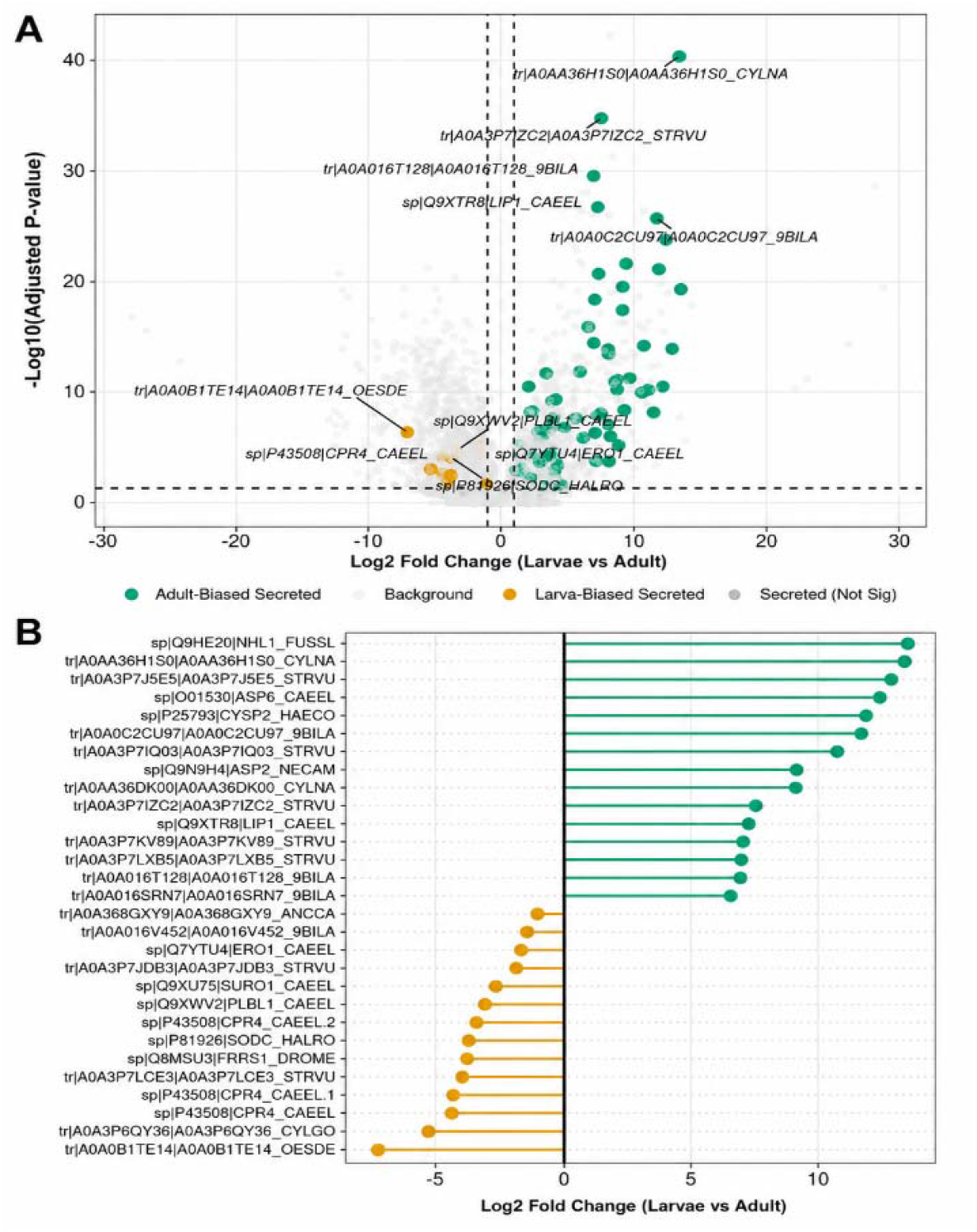
Candidate secreted proteins showed stage-associated expression patterns. Developmental expression patterns among high-confidence predicted secreted proteins. (A) Volcano plot showing predicted secreted proteins in the Larvae vs Adult differential expression contrast. Highlighted points indicate significantly stage-biased secretome candidates, with selected strongly biased proteins labeled. Dashed lines indicate fold-change and adjusted P-value thresholds. (B) Ranked lollipop plot of the strongest larva-biased and adult-biased secretome candidates by log2 fold change. Together, these panels show that the predicted S. vulgaris secretome is developmentally structured rather than uniformly expressed across parasitic stages.

Together, these results identify a focused set of candidate extracellular proteins and show that the putative secretome is developmentally structured (Figure 5B). This pattern suggests that larval and adult stages may deploy different secreted protein repertoires, consistent with their distinct host environments and biological roles.

### Stage-associated transcriptomes revealed developmental and sex-associated expression programs

Comparative transcriptome analysis showed that developmental stage was the dominant source of global expression variation across the 13 retained RNA-seq libraries. Principal component analysis and hierarchical clustering separated larval and adult samples, with additional subgroup structure among adult female, adult male, L4, ML5 female, and ML5 male libraries (Figures 6A and 6B).

**Figure 6.**
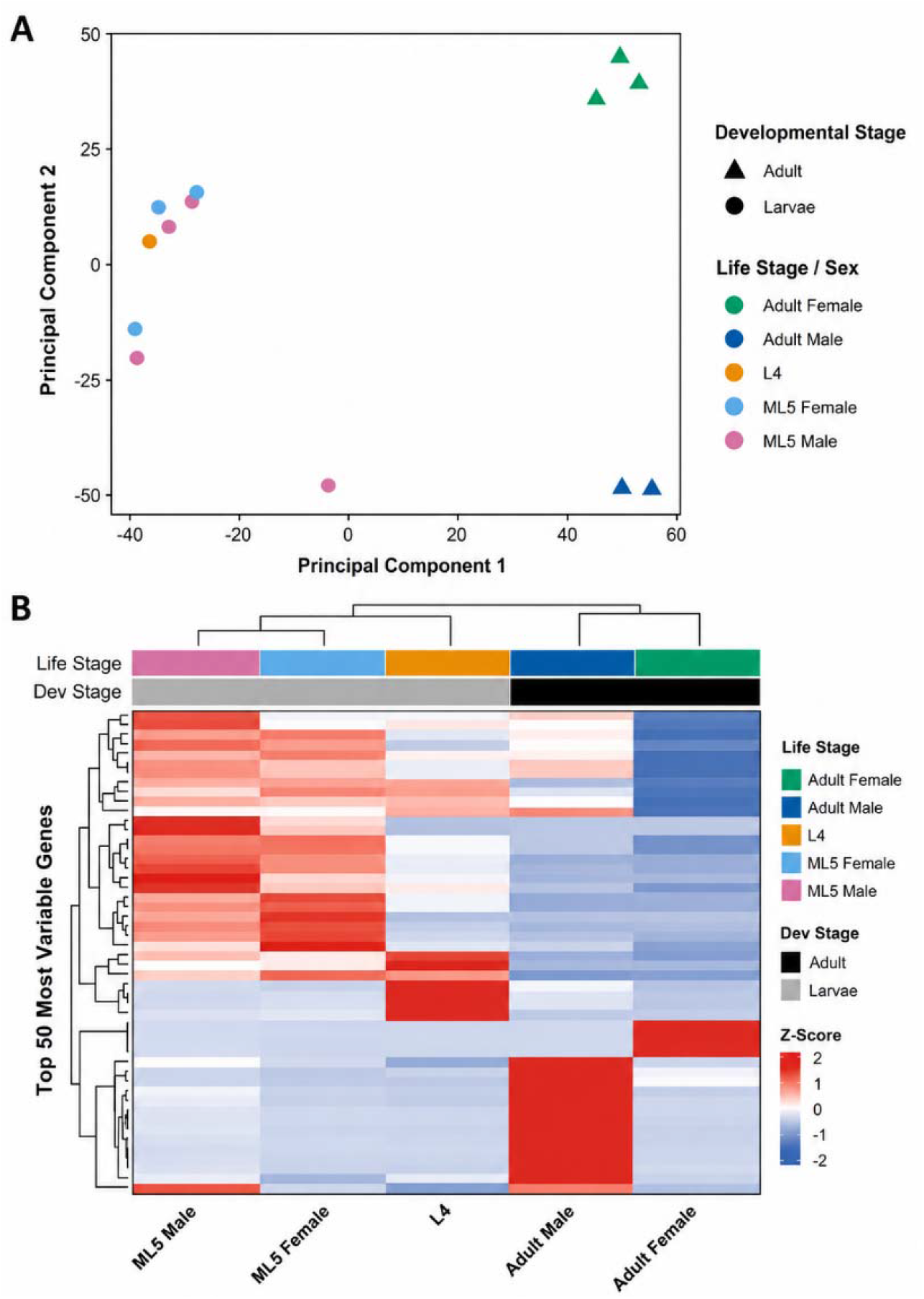
Global transcriptome structure separated larval and adult S. vulgaris samples. Global expression patterns across the 13 retained RNA-seq libraries. (A) Principal component analysis of gene-expression profiles. Samples separate primarily by developmental stage, with additional structure among libraries. (B) Hierarchical clustering heatmap of the 50 most variable genes across retained libraries. Expression values are shown as gene-level Z-scores, with sample annotations indicating life stage and developmental class. Both analyses support developmental stage as the dominant source of transcriptomic variation, with additional adult sex-associated structure.

Differential expression analyses identified widespread transcriptional divergence across developmental and sex-associated contrasts (Table 2). The strongest signals were observed between adult males and adult females, across the broader Larvae vs Adult comparison, and during the ML5 female-to-adult female transition. In contrast, only 31 genes were differentially expressed between ML5 males and ML5 females, indicating limited sex-associated transcriptional divergence during the migratory larval stage.

**Table 2.**
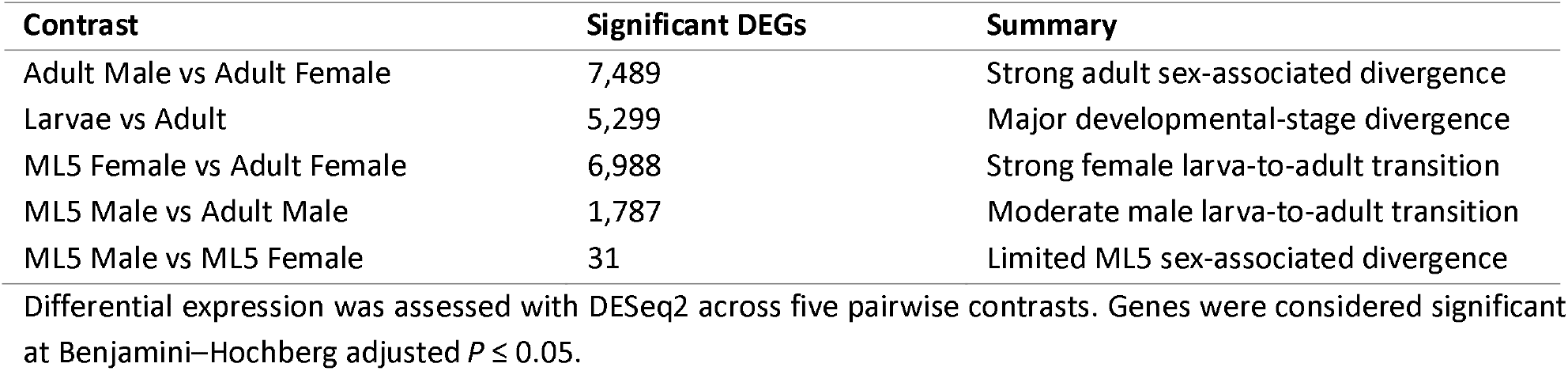
Summary of differential gene expression across developmental and sex-associated contrasts in Strongylus vulgaris.

Annotated differentially expressed genes included protease-related, collagen/cuticle-associated, cell-cycle-associated, lysozyme-like, ASP-like, and nematode-specific loci, along with many strongly responsive genes lacking informative functional names (Figures 7A and 7B). A complete differential expression summary is provided in Additional file 5.

**Figure 7.**
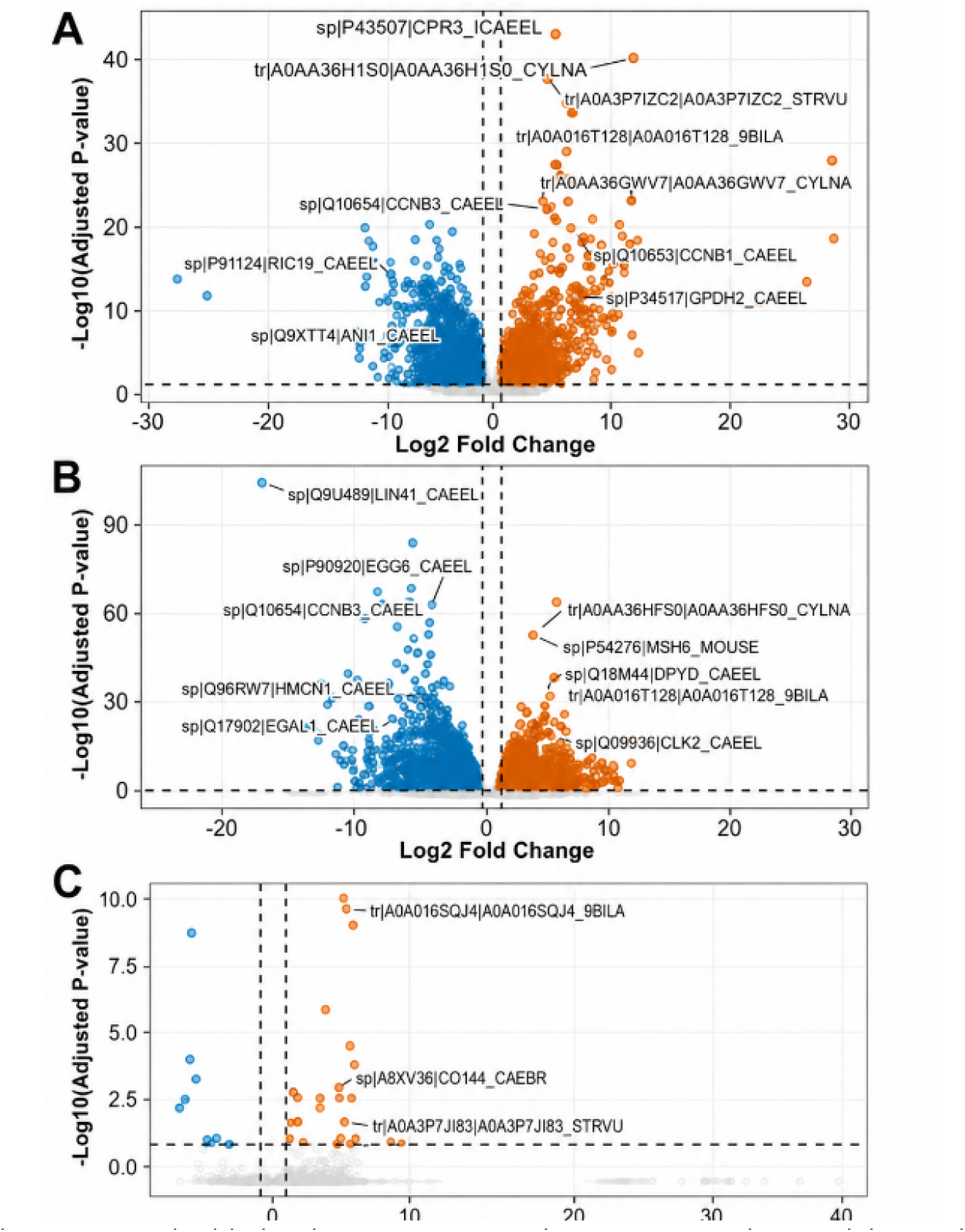
Differential expression highlighted stage-associated transcriptional remodeling.Volcano plots showing differential gene expression across major developmental contrasts. (A) Larvae vs Adult comparison, showing widespread stage-associated transcriptional divergence. (B) ML5 Female vs Adult Female comparison, highlighting transcriptional remodeling during the female migratory-larva-to-adult transition. (C) ML5 Male vs ML5 Female comparison, showing comparatively limited sex-associated differential expression during the migratory larval stage. Points represent genes, colored points indicate significantly differentially expressed loci, and selected highly responsive annotated candidates are labeled. Dashed lines indicate fold-change and adjusted P-value thresholds.

Functional enrichment analyses supported conservative interpretation of stage-associated molecular remodeling. In the Larvae vs Adult contrast, the KEGG NOD-like receptor signaling pathway label was enriched; however, the contributing genes were dominated by conserved proteolysis, stress-signaling, vesicle-trafficking, and cellular-remodeling components. This enrichment was therefore interpreted as a conserved functional module rather than evidence of a canonical vertebrate-like NOD-like receptor pathway in S. vulgaris (Additional file 6). Overall, the expression data indicate that S. vulgaris does not maintain a single uniform parasitic transcriptional profile across development. Instead, transcriptional programs shift substantially between larval and adult stages, while sex-biased expression becomes most pronounced in adults.

### Isoform diversity highlighted transcript-level regulation across developmental transitions

Isoform-level analysis identified substantial transcript diversity within the refined S. vulgaris annotation, with 24,546 transcript models quantified across the 13 RNA-seq libraries. After filtering and DEXSeq-based testing, 490 high-confidence isoform switches were identified across replicated-group contrasts. Because the L4 group was not replicated, isoform-switch testing was restricted to Adult Male vs Adult Female, ML5 Female vs Adult Female, ML5 Male vs Adult Male, and ML5 Male vs ML5 Female comparisons.

Isoform switching was most pronounced in the ML5 Female vs Adult Female contrast, which accounted for the dominant share of detected switches (Figure 8A). In contrast, isoform-level differences between ML5 males and ML5 females were comparatively limited, consistent with the weaker sex-associated transcriptional divergence observed during the migratory larval stage.

**Figure 8.**
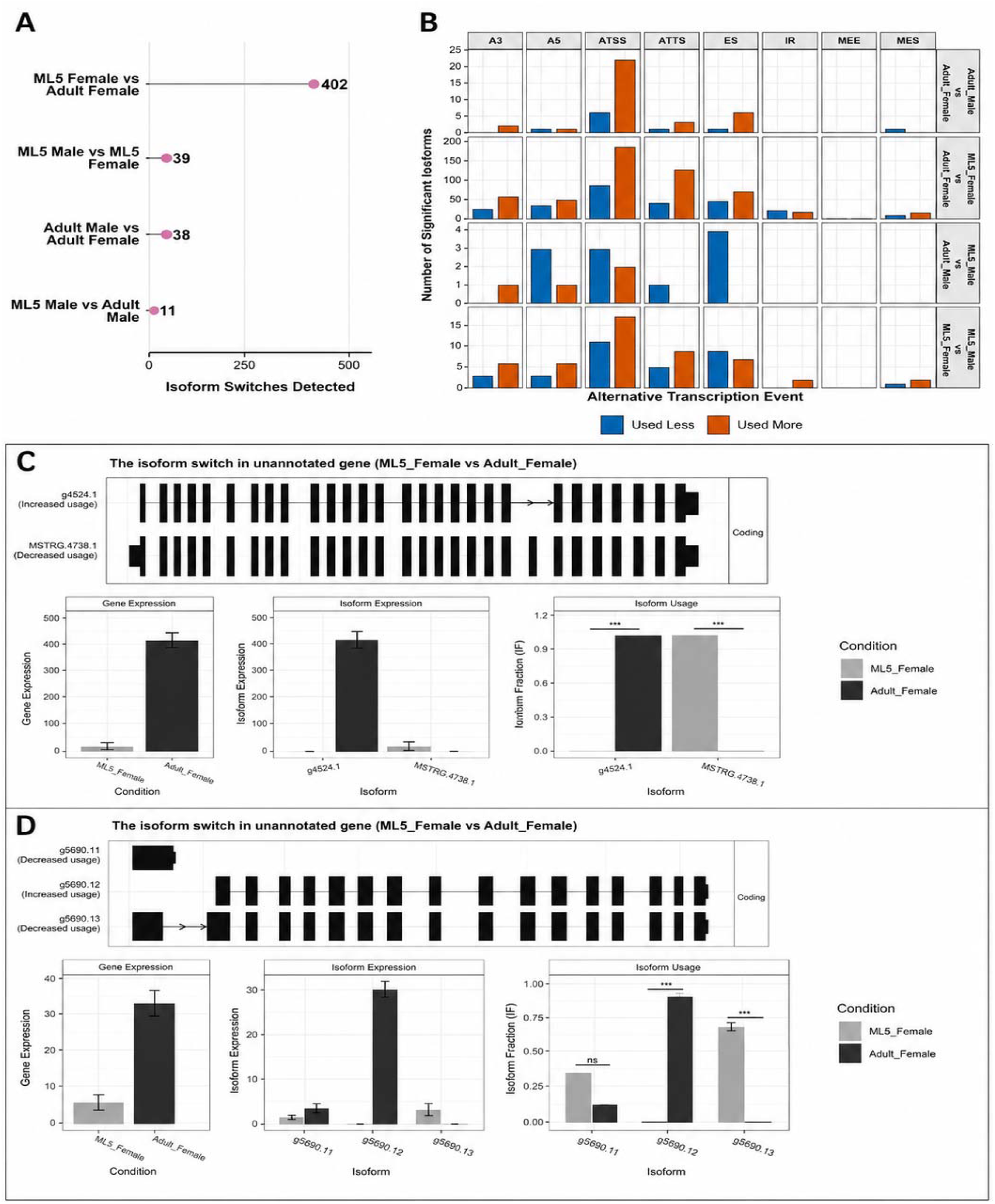
Isoform-switch analysis revealed transcript-level regulation across developmental transitions.Isoform-level analyses of transcript usage across replicated developmental and sex-associated contrasts. (A) Number of high-confidence isoform switches detected in each tested contrast. Isoform switching was most pronounced in the ML5 Female vs Adult Female comparison. (B) Alternative splicing and transcript-structure event classes contributing to significant isoform switches, including alternative transcription start-site usage, alternative transcription termination-site usage, exon skipping, alternative splice-site usage, and intron retention. Bars distinguish isoforms with increased versus decreased usage. (C) Representative isoform switch in an unannotated gene, showing reciprocal usage of g4524.t1 and MSTRG.4738.1 between ML5 female and adult female samples. (D) Representative isoform switch in g5690, showing altered usage among three transcript isoforms between ML5 female and adult female samples. Transcript models are shown above expression and isoform-usage summaries; bars represent condition-level expression or isoform fraction, with significance indicated above comparisons.

Alternative splicing analysis showed that isoform variation reflected both alternative transcript-boundary usage and splice-junction remodeling, rather than a single dominant mechanism (Figure 8B). Representative structure plots further illustrated development-associated isoform switching in unannotated loci during the ML5 female-to-adult female transition, including examples with reciprocal changes in dominant isoform usage (Figures 8C and 8D). A complete listing of high-confidence isoform switches is provided in Additional file 7.

Overall, these results show that transcript-level regulation accompanies developmental transitions in S. vulgaris, particularly during the female ML5-to-adult transition. Because many switched loci remain poorly annotated, developmentally important regulation may occur in genes not yet functionally resolved by conventional homology-based annotation.

### Comparative genomics identified lineage-specific gene-family turnover in *S. vulgaris*

Orthology inference placed S. vulgaris within a comparative framework of sampled nematodes. OrthoFinder identified 27,641 orthogroups across the dataset, including conserved shared orthogroups, strict single-copy orthologues, and S. vulgaris-specific orthogroups (Figure 9A). The rooted species tree placed S. vulgaris as sister to Cylicocyclus nassatus within the sampled equine strongylid framework (Figure 9B).

**Figure 9.**
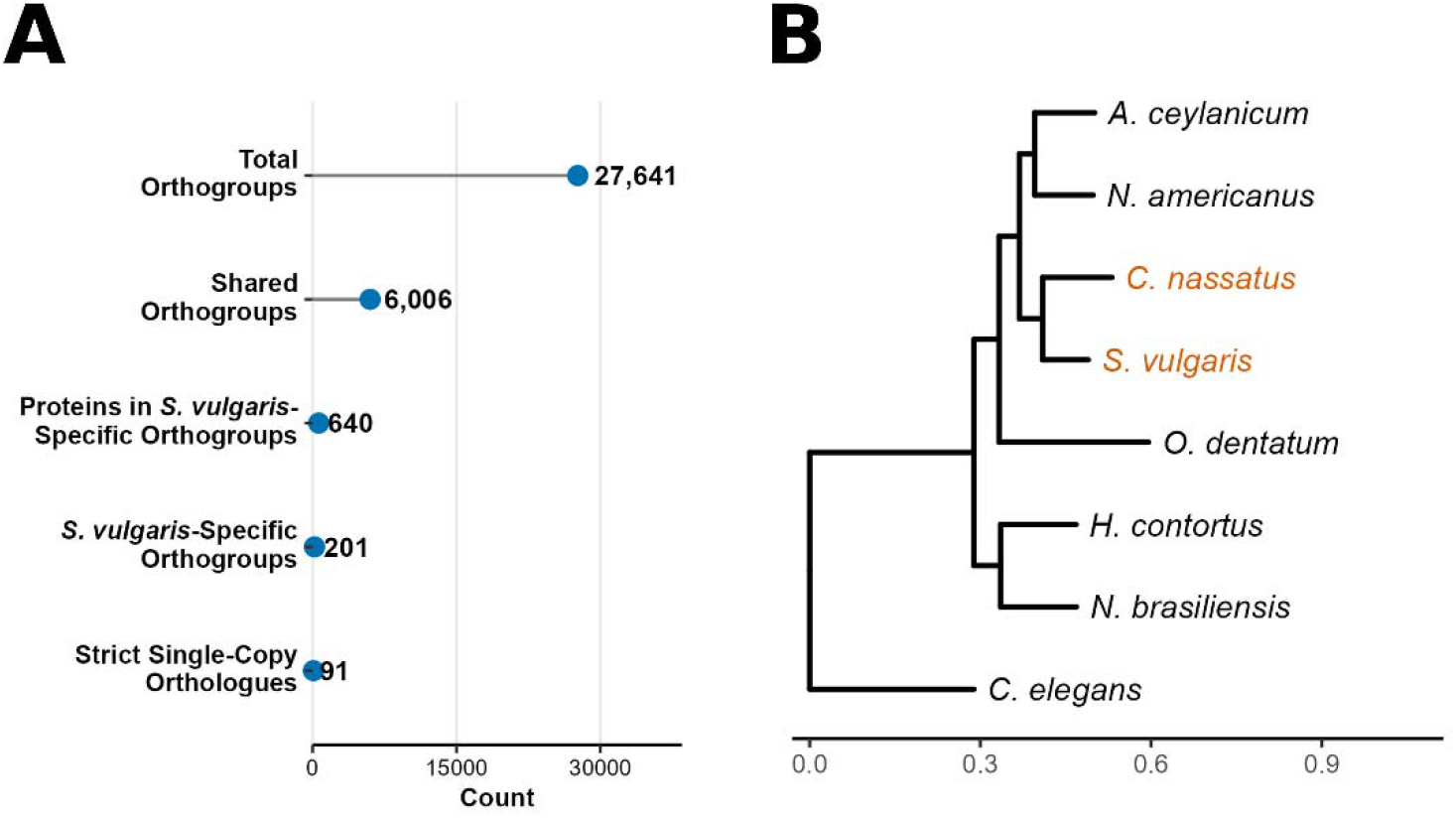
Orthology analysis placed S. vulgaris within the sampled nematode comparative framework. Orthology and phylogenetic summaries from comparative nematode analysis. (A) OrthoFinder orthogroup summary showing total orthogroups, shared orthogroups, S. vulgaris-specific proteins, S. vulgaris-specific orthogroups, and strict single-copy orthologues. (B) Rooted species tree inferred from single-copy orthologues. S. vulgaris was placed as sister to Cylicocyclus nassatus within the sampled strongylid nematodes, consistent with their shared placement among equine strongylids.

Lineage-specific orthogroup analysis identified 201 orthogroups unique to S. vulgaris, comprising 640 proteins (Additional file 8). These species-restricted families included both annotated and weakly annotated groups, indicating that the S. vulgaris proteome contains a distinct lineage-specific component in addition to its conserved nematode orthology core.

Direct comparison of orthogroup copy number between S. vulgaris and C. nassatus showed that many shared families were broadly conserved, while a subset showed marked copy-number divergence between the two equine strongylids (Figures 10A and 10B). Families with higher relative copy number in S. vulgaris included both shared expanded families and S. vulgaris-specific families absent from C. nassatus, whereas contracted families showed reduced copy number in S. vulgaris relative to C. nassatus. This pattern indicates that divergence between these sister taxa includes both gene-family presence/absence differences and quantitative shifts in copy number among shared families.

**Figure 10.**
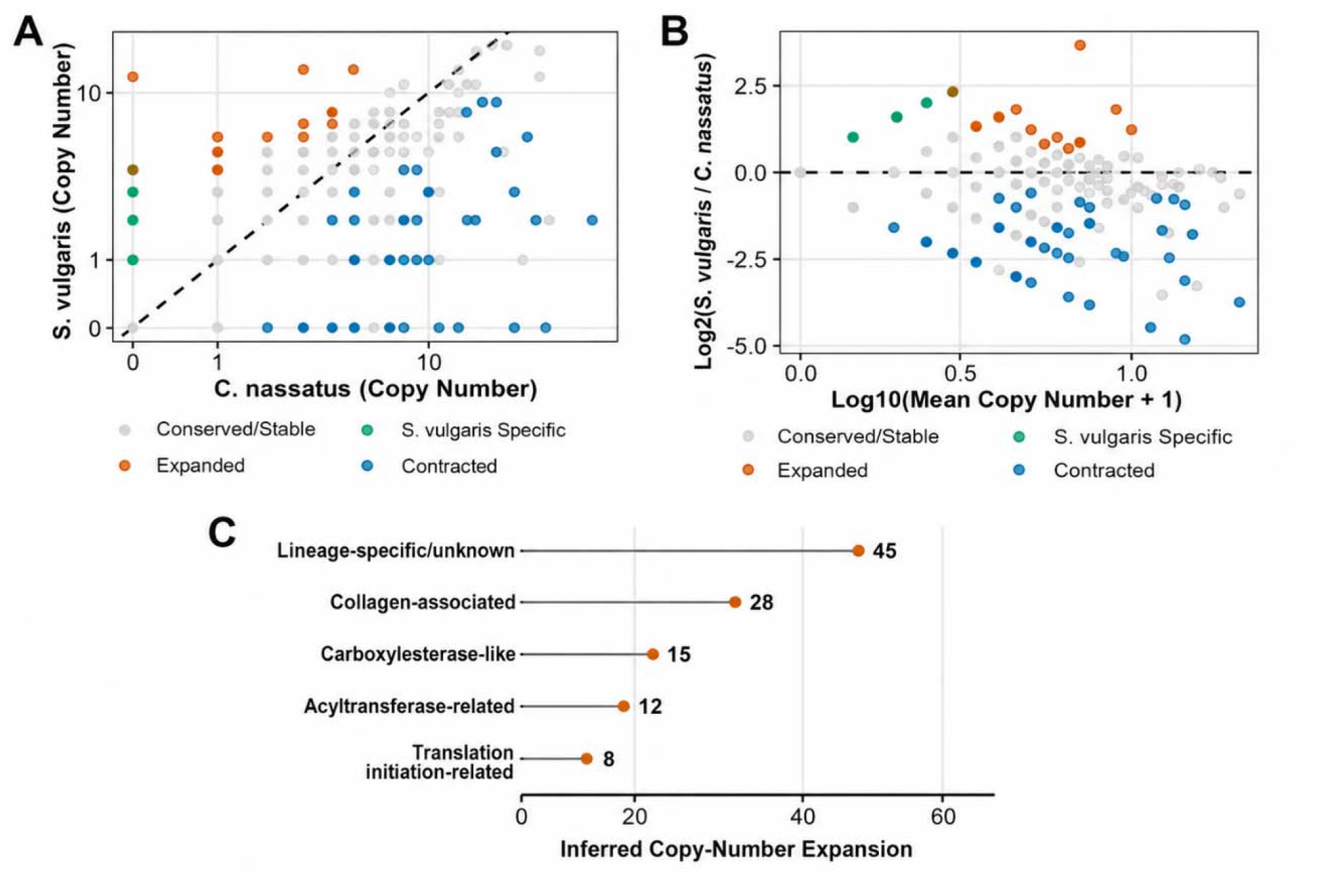
Orthogroup copy-number comparisons revealed gene-family divergence between S. vulgaris and C. nassatus. Pairwise comparison of orthogroup copy number between S. vulgaris and Cylicocyclus nassatus. (A) Raw copy-number scatterplot showing orthogroup copy numbers in S. vulgaris relative to C. nassatus. The dashed line indicates equal copy number between species. Points are classified as conserved/stable, S. vulgaris-specific, expanded in S. vulgaris, or contracted in S. vulgaris. (B) MA-style plot showing relative copy-number differences as log2(S. vulgaris/C. nassatus) against mean orthogroup copy number. Positive values indicate higher relative copy number in S. vulgaris, whereas negative values indicate lower relative copy number. (C) Major functional categories among expanded S. vulgaris gene families after filtering repeat-associated and extreme copy-number artifacts. Expanded categories included lineage-specific or unknown families, collagen-associated proteins, carboxylesterase-like proteins, acyltransferase-related proteins, and translation-initiation-related families.

CAFE5 gene-family evolution analysis further supported substantial lineage-specific turnover in S. vulgaris. After filtering repeat-associated and extreme copy-number artifacts, 48 families were classified as significantly expanded and 250 as significantly contracted in the S. vulgaris lineage (Additional file 9). Prominent expanded categories included lineage-specific or weakly annotated families, collagen-associated families, carboxylesterase-like proteins, acyltransferase-related proteins, and translation-initiation-related families (Figure 10C).

Together, these comparative analyses show that S. vulgaris retains a conserved nematode orthology core while also exhibiting species-specific orthogroups and substantial gene-family copy-number turnover. The expansion of weakly annotated and lineage-restricted families suggests that part of the S. vulgaris coding repertoire may represent novel or highly diverged gene families that warrant future functional investigation.

### Structure-guided annotation prioritized migratory-stage secreted proteins with candidate host-interface functions

Structure-guided annotation was applied to 1,327 proteins that remained unknown or weakly annotated after conventional homology-, domain-, and pathway-based annotation. These candidates represented 1,137 genes and 1,325 transcript models and were retained for structural analysis because they lacked informative Swiss-Prot annotation or strong conventional functional support. AlphaFold2, Foldseek, DeepFRI, COFACTOR, and DALI analyses improved interpretability for a subset of these proteins, but most candidates were retained conservatively as fold-level or structurally supported predictions rather than definitive functional assignments.

A subset of 202 structurally prioritized proteins was also predicted to belong to the secretome. Of these, 132 were upregulated in migratory-stage comparisons, suggesting that part of the weakly annotated secretome is enriched during tissue-associated larval development. Structurally interpretable candidates included VWA-domain, CAP/SCP/TAPS-like, cysteine-rich, Plasminogen/Apple/Nematode (PAN) domain-containing proteins, DNase II-like, thrombospondin-domain, Stichodactyla helianthus potassium channel toxin (ShK) domain-containing proteins, and lipid-binding-like proteins, consistent with potential extracellular, adhesive, proteolytic, or host-interface roles (Figure 11; Additional file 10).

**Figure 11.**
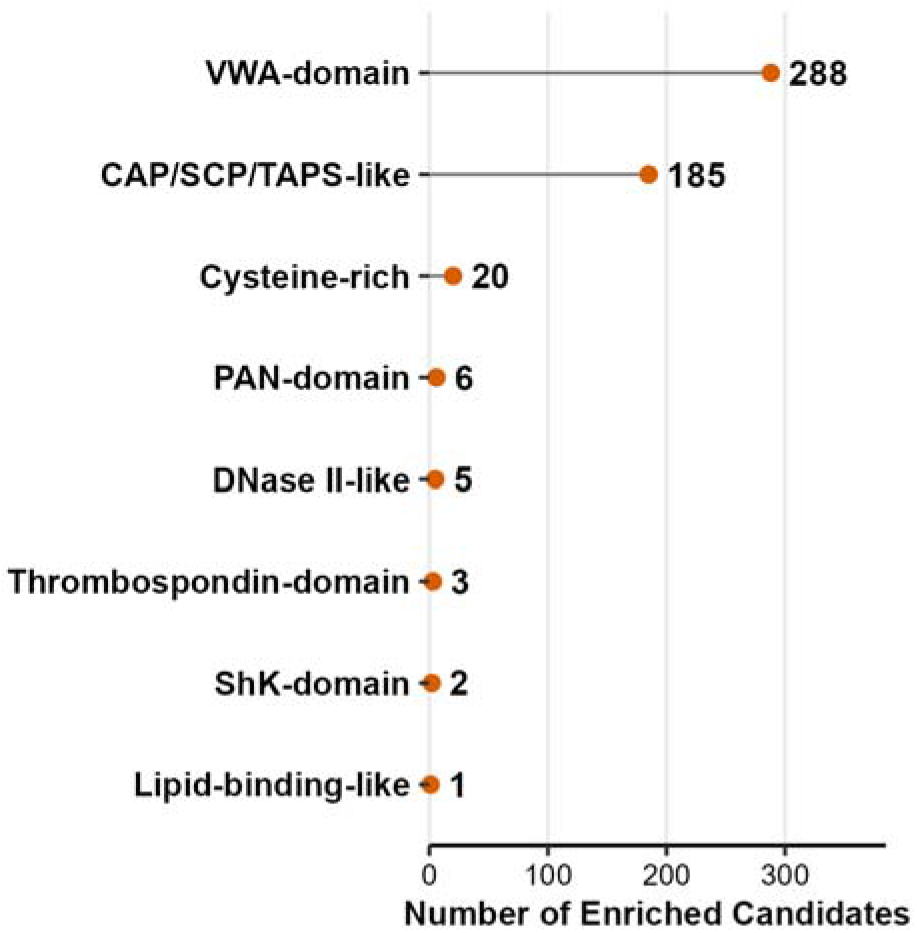
Structure-guided annotation identified major classes of enriched weakly annotated candidate proteins. Structural and domain-associated classes among enriched weakly annotated candidate proteins prioritized by structure-guided annotation. VWA-domain and CAP/SCP/TAPS-like candidates represented the largest recognizable groups, followed by cysteine-rich, PAN-domain, DNase II-like, thrombospondin-domain, ShK-domain, and lipid-binding-like candidates. Counts represent candidate class assignments and should be interpreted as fold- or domain-level prioritizations rather than definitive biochemical functional assignments.

Several candidate classes were especially notable. VWA-domain and CAP/SCP/TAPS-like proteins represented the largest recognizable structural categories among enriched candidates, while smaller groups included cysteine-rich, PAN-domain, DNase II-like, thrombospondin-domain, ShK-domain, and lipid-binding-like proteins. These annotations were interpreted at the fold or domain level, as structural similarity alone did not support assignment of specific ligands, host targets, or biochemical activities.

A smaller prioritized set of 19 proteins met combined criteria for predicted secretion, migratory-stage differential expression, limited conventional annotation, and potential relevance to host-interface or developmental biology. Within this set, COFACTOR identified SVUL_124530.1.p1 as the strongest functional candidate, supporting a conservative annotation as a putative secreted peptidase-like protein with procathepsin B-like structural similarity. DALI comparisons provided additional structural support for selected candidates, including alpha-helical or coiled-coil-like peptides, cysteine-rich secreted proteins, repeat-rich secreted proteins, and an apolipoprotein-domain candidate, but most remained functionally unresolved.

Overall, structure-guided annotation revealed that S. vulgaris expresses a diverse repertoire of migratory-stage-enriched candidate secreted proteins that are poorly resolved by conventional homology-based annotation. Their secretion signals, migratory-stage enrichment, and structural features make them strong candidates for future studies of tissue migration, host-interface biology, and stage-specific parasite development.

### Conservative HGT screening identified a small set of expressed candidate loci

A conservative alienness-based screen was used to identify putative horizontal gene transfer candidates in the S. vulgaris predicted proteome. Initial DIAMOND searches and Alien Index scoring identified 316 proteins with non-metazoan similarity patterns, of which 84 met the strict Alien Index threshold of ≥45. After additional filtering for RNA-seq support, scaffold context, fragmentary sequence, excessive amino acid identity to non-metazoan matches, and database-artifact annotations, 17 proteins were retained as strict putative HGT candidates. These represented 16 unique transcript loci because two retained proteins were isoforms from the same locus (Additional file 11; Table 3).

**Table 3.**
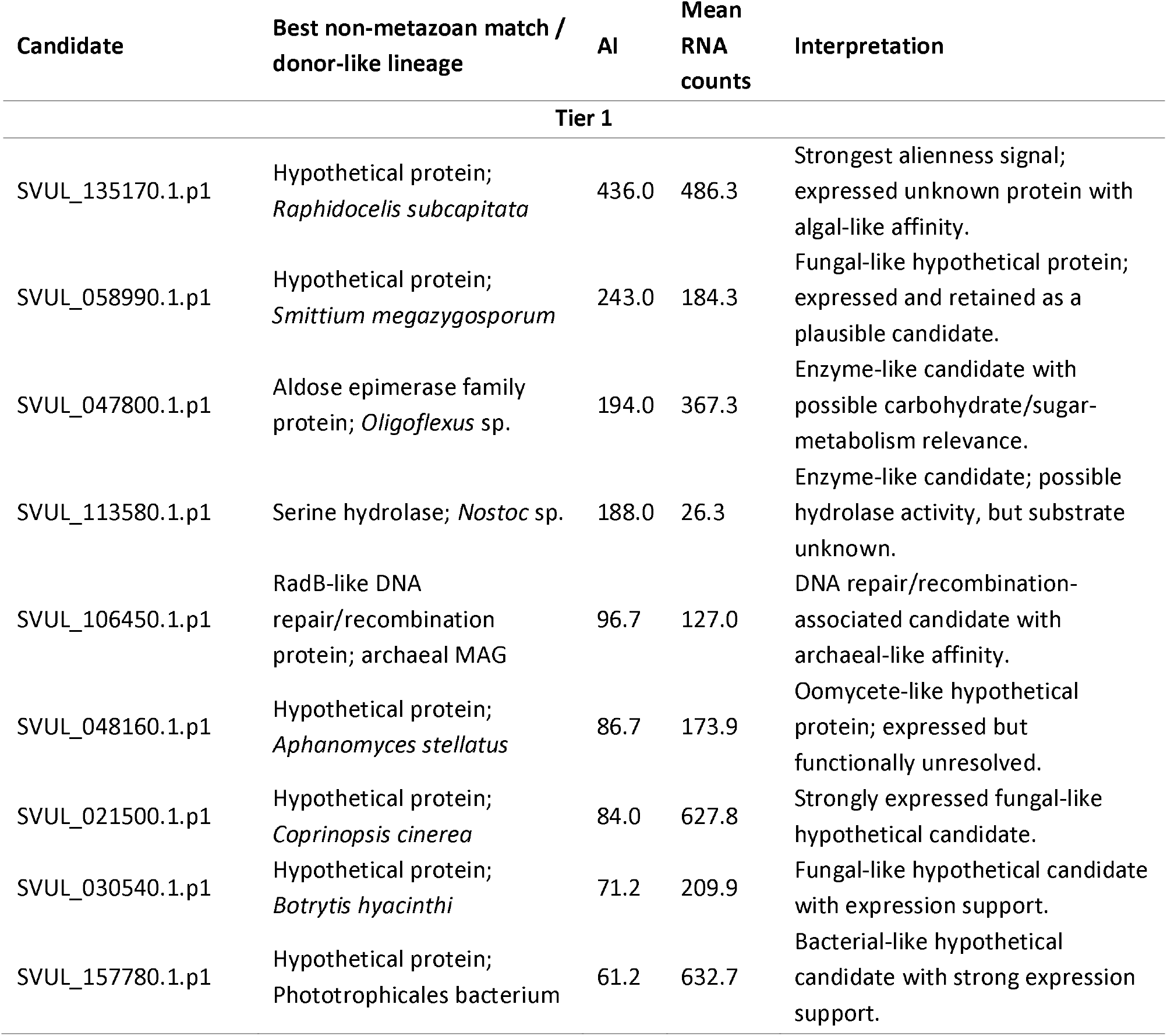
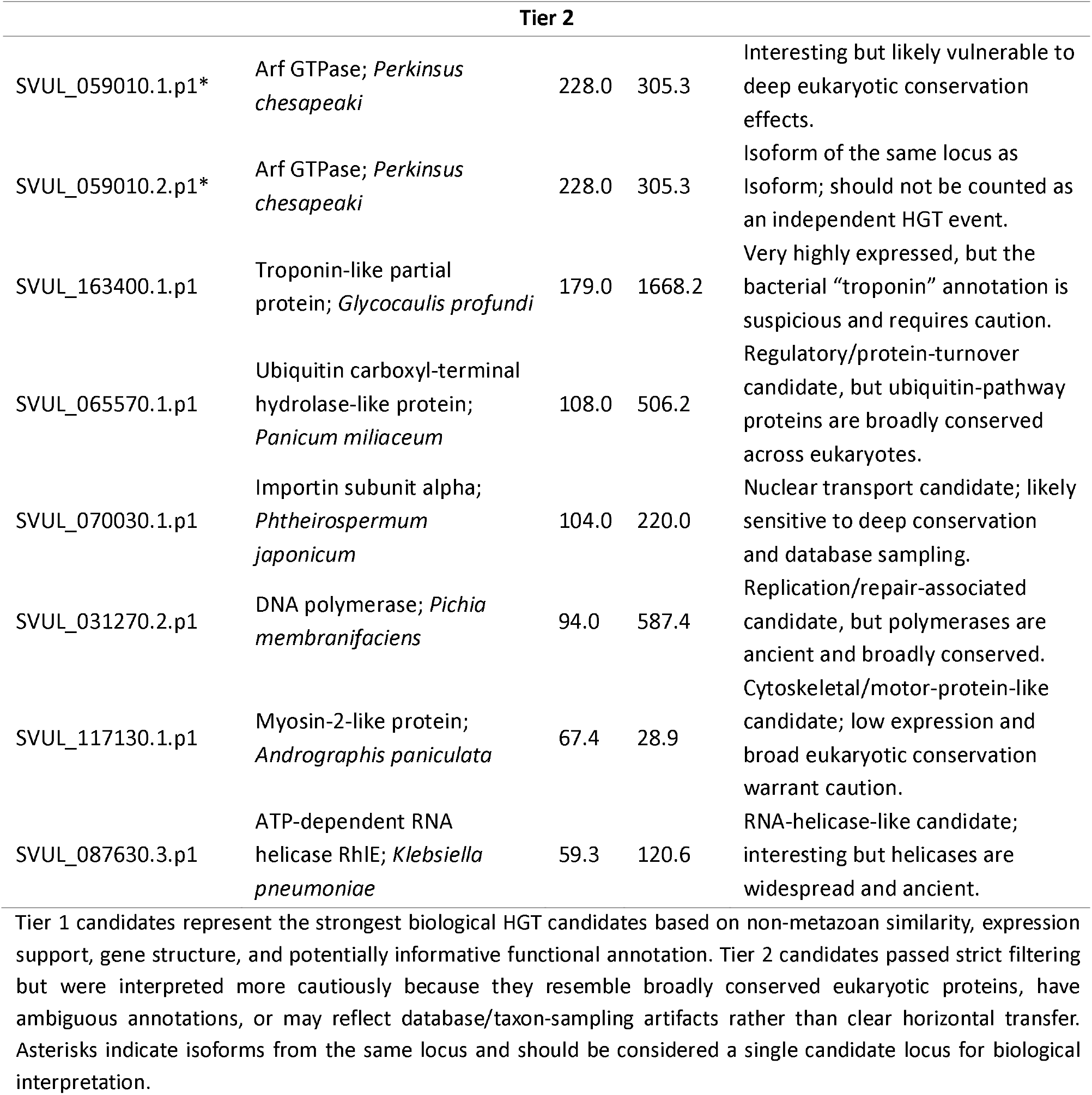
Tiered interpretation of 17 strict putative HGT candidates in Strongylus vulgaris.

The retained candidates had Alien Index values ranging from 59.3 to 436.0 and were supported by RNA-seq expression. All were annotated as complete, multi-exonic gene models, with scaffold placement and transcript support arguing against simple unexpressed microbial contamination. However, these features do not independently prove horizontal transfer and were interpreted as prioritization evidence rather than definitive validation.

Best non-metazoan matches spanned bacterial or cyanobacterial, fungal, protist/algal or stramenopile-like, plant, and archaeal lineages. Several candidates matched hypothetical or poorly characterized proteins, whereas others resembled broadly conserved eukaryotic proteins, including GTPase-, importin-, ubiquitin hydrolase-, myosin-, polymerase-, and helicase-like proteins. Candidates resembling broadly conserved eukaryotic proteins were interpreted with particular caution because deep conservation, uneven taxon sampling, or database imbalance can produce HGT-like similarity patterns.

Composition-based analyses supported this conservative interpretation. Retained candidates showed some GC-content and codon-usage structure but remained broadly embedded within the native coding-gene background (Figures 12A and 12B). Expression-versus-Alien Index comparisons further showed that retained candidates were expressed across a wide range of support values rather than being driven solely by low-expression artifacts (Figure 12C).

**Figure 12.**
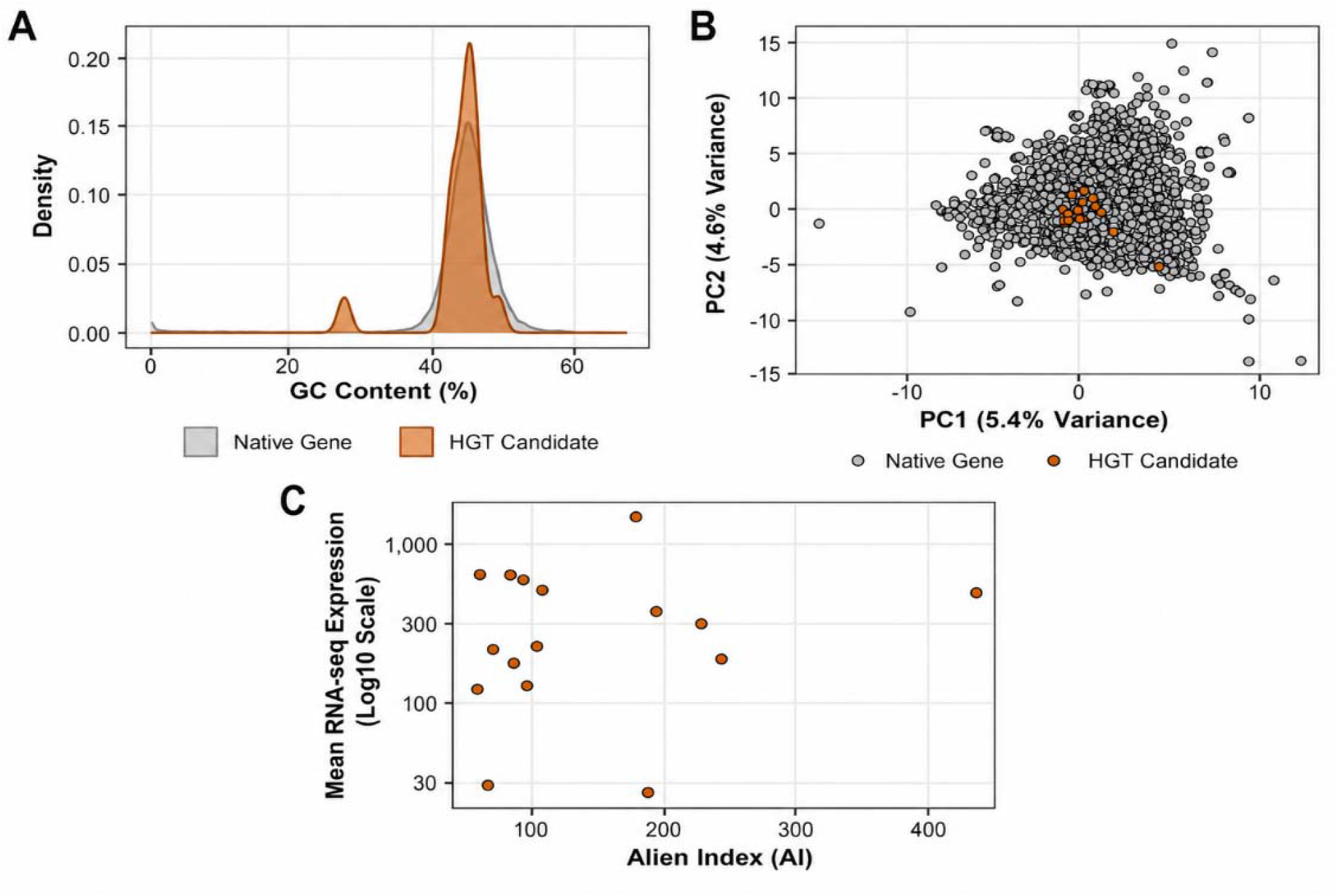
Composition and expression analyses supported conservative prioritization of putative HGT candidates. Supporting analyses for strict putative horizontal gene transfer candidates retained after Alien Index screening and conservative filtering. (A) GC-content distribution of retained HGT candidates compared with the native coding-gene background. HGT candidates showed partial compositional distinction but overlapped broadly with native genes. (B) Codon-usage principal component analysis comparing retained HGT candidates with native coding genes. Candidate genes occupied a restricted region of codon-usage space but remained embedded within the broader native distribution. (C) Relationship between Alien Index score and mean RNA-seq expression for retained candidates. Retained candidates showed expression support across a range of Alien Index values, supporting their prioritization as expressed candidate loci rather than unexpressed contaminant sequences.

Together, Alien Index scoring, expression support, intron-bearing gene structure, scaffold context, composition analysis, and preliminary phylogenetic screening identified a small set of high-priority putative HGT candidates in S. vulgaris. These candidates remain best interpreted as putative transfers requiring additional phylogenetic and experimental validation.

### Resistance-associated homologs provide a baseline for future surveillance

To establish a genomic baseline for future resistance surveillance, 43 curated resistance-associated query sequences from C. elegans, cyathostomins, and other parasitic nematodes were screened against the S. vulgaris genome and representative proteome. Homologs were recovered for all 43 queries, including 34 high-confidence and 9 moderate-confidence assignments. Because multiple queries often matched the same S. vulgaris protein, these hits collapsed to 18 nonredundant S. vulgaris homologs with recoverable genomic coordinates (Additional file 12).

Detected homologs included β-tubulin-related proteins, P-glycoprotein/ABC-transporter-associated proteins, a levamisole receptor-associated lev-1-like homolog, a Dyf7-like homolog, and additional resistance-associated proteins previously identified in strongylid nematodes. These results indicate that major surveyed resistance-associated gene families are represented in the S. vulgaris genome, although homolog detection alone does not imply phenotypic resistance.

The β-tubulin orthogroup provided the clearest marker-level result. Four S. vulgaris β-tubulin homologs were present in the targeted orthogroup tree, and all retained susceptible-state residues at the canonical benzimidazole-associated positions F167, E198, and F200. No resistant-state substitutions were detected at these sites.

Copy-number and phylogenetic analyses supported a conservative interpretation. Targeted phylogenies generally placed S. vulgaris homologs within broader nematode orthogroups, often near strongylid or cyathostomin homologs, supporting their classification as conserved homologs rather than isolated assembly artifacts. No targeted resistance-associated family showed evidence of clear S. vulgaris-specific expansion.

Expression-based interpretation was limited. Although all resistance-associated homologs were supported by Iso-Seq evidence, RNA-seq overlays did not identify a robust stage-biased expression pattern among these loci. Expression data were therefore not used to infer resistance-associated stage regulation.

Together, these analyses provide a baseline catalog of resistance-associated homologs in S. vulgaris. The surveyed loci are structurally recoverable and evolutionarily interpretable, but the results do not provide evidence for phenotypic resistance, canonical β-tubulin resistance substitutions, or expansion of resistance-associated gene families. Instead, this analysis identifies candidate loci for future population-level surveillance, comparative analysis, and targeted validation if resistant or treatment-selected S. vulgaris material becomes available.

## Discussion

This study establishes a substantially improved genomic and transcriptomic framework for S. vulgaris, a clinically important equine strongyle whose molecular biology has remained underdeveloped despite its well-recognized pathogenicity [11]. The new PacBio HiFi assembly substantially improves the prior short-read draft genome reported by the International Helminth Genomes Consortium, increasing assembly continuity and conserved-gene recovery while providing a stronger foundation for transcript-supported annotation, functional interpretation, comparative genomics, and candidate-gene discovery [16]. More importantly, these resources allow S. vulgaris biology to be examined across multiple molecular layers for the first time: genome architecture, transcript structure, protein annotation, secretome prediction, stage-associated expression, isoform diversity, gene-family evolution, putative HGT candidates, and resistance-associated homologs.

### Genome architecture and genome size in *S. vulgaris*

The S. vulgaris genome appears to be shaped more strongly by repetitive sequence and noncoding architecture than by broad expansion of the conventional protein-coding gene repertoire. The large repetitive fraction, particularly the dominance of unclassified repeats, suggests that genome size reflects substantial repeat accumulation and repeat divergence. The high proportion of unclassified repetitive sequence is especially informative because it indicates that much of the repetitive compartment is either lineage-specific, highly diverged, or insufficiently represented in current repeat libraries.

This interpretation distinguishes genome architecture from gene-family expansion. Although comparative analyses identified lineage-specific gene-family turnover, the annotated gene set and filtered comparative proteome do not support a model of genome inflation driven primarily by wholesale protein-coding expansion. Instead, the data suggest that repetitive sequence, isoform richness, and selected gene-family changes jointly shape the S. vulgaris genome. This distinction is important because repeat-rich architecture can influence genome organization, annotation complexity, recombination, regulatory evolution, and comparative inference even when gene number remains within the expected range for parasitic nematodes.

The assembly also provides a basis for future studies of structural genome evolution. Because the present assembly is not chromosome-scale, it cannot fully resolve long-range repeat organization, segmental duplications, synteny, or chromosomal rearrangements. These features will be important for determining whether genome architecture in S. vulgaris reflects gradual repeat accumulation, lineage-specific repeat activity, localized duplication, or broader chromosomal restructuring.

### Evolutionary divergence among equine strongylids

The comparative genomic results indicate that S. vulgaris retains a conserved nematode orthology core while also showing lineage-specific components that distinguish it from sampled relatives. This pattern is consistent with a model in which divergence between large and small strongyles is not driven by wholesale replacement of the conserved gene complement, but by differential retention, expansion, contraction, and possible innovation within selected gene families.

The comparison with Cylicocyclus nassatus is particularly useful because these taxa are evolutionarily close enough for shared orthology to be informative but biologically distinct in tissue tropism and pathogenic strategy. Cyathostomin disease is associated primarily with larval development in the intestinal wall, whereas S. vulgaris undergoes arterial migration and is associated with vascular pathology [3–11]. The observed gene-family differences therefore provide candidate molecular axes along which these divergent life histories may have evolved.

Expanded or lineage-associated categories such as collagen-related proteins, carboxylesterase-like proteins, acyltransferase-related proteins, and weakly annotated families may reflect differences in cuticle biology, developmental remodeling, metabolism, detoxification, or host-interface biology. These interpretations remain provisional because copy-number change alone does not establish adaptive function. However, they identify a biologically coherent set of candidate families for future comparative work across additional large strongyles and cyathostomins.

The current taxon sampling limits how far these evolutionary conclusions can be extended. Some features identified here may be specific to S. vulgaris, while others may be shared among large strongyles or distributed more broadly across Strongylida. Broader comparative sampling, particularly with chromosome-scale assemblies, will be needed to distinguish species-specific changes from traits associated with arterial migration, large-strongyle biology, or strongylid diversification more generally.

### Resistance-associated homologs as a baseline for surveillance

The resistance-associated homolog survey is best interpreted as a genomic baseline rather than as a resistance study. The recovery of homologs corresponding to major nematode drug-response-associated gene families shows that these loci are structurally represented in S. vulgaris, but it does not imply resistance. This distinction is essential because homolog presence, even at loci associated with drug response in other nematodes, is not equivalent to phenotypic resistance or treatment-selected variation.

The absence of canonical β-tubulin benzimidazole-resistance substitutions in the reference genome is therefore most appropriately interpreted as a baseline marker-state observation. Similarly, the lack of obvious S. vulgaris-specific expansion among surveyed resistance-associated families argues against major lineage-specific amplification of these loci in this assembly, but it does not address population-level variation or treatment response. Those questions require resequencing of field populations, explicit comparison across epidemiological settings, and, where possible, phenotypic or treatment-outcome data.

This baseline is valuable because resistance is a major concern in equine cyathostomins and other parasitic nematodes, whereas S. vulgaris resistance has not been documented in the same way and remains difficult to evaluate species-specifically [1,11,42–45,67]. The genomic coordinates, orthogroup placement, and marker-state information generated here provide a reference framework for future surveillance if changing parasite-control practices alter S. vulgaris prevalence, transmission, or selection pressure.

### Developmental regulation at the gene and isoform levels

The transcriptomic results support development as the dominant organizing axis of S. vulgaris molecular biology. Rather than maintaining a uniform parasitic expression program across life stages, S. vulgaris appears to remodel gene expression substantially between migratory larval and adult stages. This pattern is biologically consistent with the parasite’s transition from tissue-associated and arterial environments to intestinal adult habitats, where reproductive specialization becomes a major feature of adult biology [6,8,11].

The limited transcriptional divergence between ML5 males and females, contrasted with strong adult sex-associated divergence, suggests that sex-specific programs become most pronounced after the migratory stage. During migration, shared developmental and host-interface requirements may dominate the transcriptome of both sexes. After maturation, reproductive differentiation likely becomes a stronger driver of gene expression. This provides a developmental interpretation that connects the RNA-seq patterns to the parasite’s life history rather than simply cataloging differentially expressed genes.

Isoform switching adds an additional regulatory dimension to this model. The concentration of isoform switches during the female ML5-to-adult transition suggests that maturation involves not only changes in transcript abundance but also changes in transcript structure. Such remodeling could alter coding potential, untranslated regions, domain composition, transcript stability, or nonsense-mediated decay sensitivity. Developmentally regulated alternative splicing and nonsense-mediated decay have been described in C. elegans [23,24], and the present data suggest that analogous mechanisms may be relevant in a parasitic strongylid.

The biological importance of this finding is heightened by the poor annotation of many switched loci. If developmentally regulated isoforms occur in genes that lack informative homology, then a substantial component of S. vulgaris developmental regulation may reside in lineage-specific or highly diverged gene space. This underscores the importance of retaining isoform-aware annotations in non-model parasite genomes, where collapsing to a single representative transcript can obscure regulatory variation that may be biologically meaningful.

### Pathway remodeling and candidate host-interface biology

Functional enrichment and pathway analyses support broad developmental remodeling, but pathway labels must be interpreted through the contributing genes rather than accepted at face value. This is particularly important for KEGG terms derived from vertebrate immune or disease contexts. In S. vulgaris, such labels are more appropriately treated as indicators of conserved molecular modules involving proteolysis, stress signaling, vesicle trafficking, cellular remodeling, or signal transduction, rather than as evidence of vertebrate-like pathways in a nematode parasite.

The predicted secretome provides a more direct framework for candidate host-interface biology. The developmental structuring of secreted candidates suggests that migratory larvae and intestinal adults deploy different extracellular repertoires. This is consistent with their distinct biological environments: migratory stages encounter host tissues, vascular structures, immune effectors, and remodeling environments, whereas adults occupy the intestinal lumen and mucosal surface. Thus, the secretome results support the broader conclusion that extracellular deployment in S. vulgaris is stage-associated rather than constitutive.

Structure-guided annotation was especially useful for interpreting weakly annotated secreted candidates. Several migratory-stage-enriched candidates belonged to protein classes with plausible extracellular or host-interface roles, including CAP/SCP/TAPS-like proteins, VWA-domain proteins, cysteine-rich proteins, PAN-domain proteins, DNase II-like proteins, thrombospondin-domain proteins, ShK-domain proteins, lipid-binding-like proteins, and peptidase-like proteins. These classes are consistent with known themes in helminth and broader parasite biology, where extracellular proteins can mediate adhesion, proteolysis, lipid interaction, immune modulation, and host-interface remodeling [26–31,36,37].

CAP/SCP/TAPS-like candidates are particularly notable because venom allergen-like proteins are prominent in helminth secretomes and have been implicated in host interaction across animal-and plant-parasitic systems [29,65,66]. DNase II-like candidates are also biologically plausible given evidence that hookworm larvae can secrete DNase II to degrade neutrophil extracellular traps [31]. These parallels should be treated as hypothesis-generating rather than functional equivalence. The present data prioritize candidates but do not demonstrate secretion in vivo, biochemical activity, host binding, or immunological effect.

The central implication is that S. vulgaris contains a developmentally regulated candidate secretome enriched for weakly annotated proteins that may participate in migration, tissue interaction, immune exposure, or stage-specific host adaptation. These candidates now provide a rational basis for proteomic validation and functional testing.

### Putative horizontal gene transfer and genomic novelty

The HGT screen provides evidence for a small set of candidate loci with non-metazoan similarity patterns, but these candidates should remain explicitly provisional. In plant-parasitic nematodes, HGT has contributed to host invasion and feeding biology, particularly through genes involved in cell-wall modification and host-derived carbohydrate metabolism [32,33,41]. Comparable evidence in animal-parasitic strongylids is more limited, and therefore the S. vulgaris candidates require conservative interpretation.

The retained candidates are biologically more credible than simple database hits because they have expression support, scaffold context, and intron-bearing gene models. These features reduce the likelihood that they represent unexpressed microbial contamination. However, several candidates resemble broadly conserved eukaryotic proteins, and composition-based analyses did not clearly separate the candidate set from the native coding background. These observations allow multiple explanations, including ancient acquisition, deep conservation, uneven taxon sampling, differential gene loss, or database imbalance.

The HGT analysis therefore contributes a candidate framework rather than a definitive evolutionary claim. The strongest candidates, particularly expressed hypothetical or enzyme-like loci with high alienness scores, warrant deeper phylogenetic sampling and functional assessment. At present, they should be discussed as possible contributors to genomic novelty, not as confirmed adaptive acquisitions or established mechanisms of host adaptation.

### Integrated model and future directions

Taken together, these analyses support a model in which S. vulgaris biology is organized by the interaction of genome architecture, developmental regulation, and lineage-specific evolution. The genome is repeat-rich and contains a substantial unclassified repetitive compartment. The coding repertoire retains a conserved nematode core while also including species-specific orthogroups and selected gene-family shifts. Developmental stage structures gene expression, isoform usage, and predicted secretome deployment. The migratory-stage secretome contains weakly annotated proteins with plausible extracellular roles, and putative HGT candidates provide a limited but testable set of loci for future evolutionary investigation.

This integrated framework is consistent with the parasite’s complex life cycle, which requires intestinal invasion, arterial migration, prolonged intravascular development, return to the intestine, sexual maturation, and reproduction [6,8,11]. The molecular data suggest that these transitions are accompanied by coordinated regulatory and extracellular remodeling rather than by a single generalized parasitic program.

Several limitations remain. The assembly is not chromosome-scale, limiting interpretation of synteny, long-range repeat structure, segmental duplication, and chromosomal organization. The primary assembly represents a single individual, so population-level variation is not captured. Transcriptomic sampling was constrained by parasite availability, and whole-organism RNA-seq cannot resolve tissue- or cell-type-specific regulation. Finally, secretome prediction, structural annotation, pathway enrichment, HGT inference, and resistance-homolog screening are computational and hypothesis-generating; each identifies candidates but does not independently establish biological function.

Future work should prioritize chromosome-scale assembly, expanded developmental sampling, tissue- or cell-resolved transcriptomics, proteomic validation of secreted products, biochemical and host-interaction assays for prioritized structure-supported candidates, and population-level resequencing across epidemiologically diverse isolates. These approaches will be necessary to determine which candidate genes, isoforms, secreted proteins, gene-family shifts, or putative HGT loci contribute directly to migration, vascular persistence, host interaction, developmental regulation, and parasite-control-relevant variation.

## Conclusions

Overall, this study moves S. vulgaris beyond a primarily pathological and life-cycle-defined organism into a system that can be studied through integrated genomics, transcriptomics, structural annotation, and comparative evolution. It is a developmentally dynamic, repeat-rich, transcript-structurally complex strongylid whose molecular biology appears to be organized around stage-specific deployment of conserved and lineage-specific gene repertoires. This work provides the first comprehensive molecular framework for understanding how S. vulgaris develops, migrates, persists, and causes disease in the horse.

## Declarations

## Ethics approval and consent to participate

University of Kentucky IACUC protocol #2021-3879.

## Consent for publication

Not applicable.

## Availability of data and materials

The genomic and transcriptomic datasets generated and/or analyzed during the current study are available in the National Center for Biotechnology Information (NCBI) under BioProject PRJNA1492748.

## Authors contributions

NER: Conception, study design, data acquisition, analysis, and interpretation, and drafting and revision of manuscript. BR: Data analysis and revision of manuscript. KL: Data analysis and revision of manuscript. EH: Study design, acquisition of data, and revision of manuscript. DH: Study design and revision of manuscript. TK: Conception, study design, data acquisition, and revision of manuscript. MLS: Study design, acquisition of data, revision of manuscript. MKN: Conception, study design, acquisition of data, revision of manuscript, supervision.

## Acknowledgements

The authors extend sincere gratitude to Dr. Stephen Doyle for his guidance, encouragement, and support throughout this endeavor. We are also deeply grateful to Research Analyst Holli Gravatte, the staff of the University of Kentucky Veterinary Diagnostic Laboratory for their patience and assistance, and the many undergraduate researchers whose help with herd care and specimen collection made this work possible.

## Additional file descriptions

**Additional file 1**

File format: Portable Network Graphics image (.png) Title of data: Additional file 1. Bioinformatic and statistical analysis pipeline for Strongylus vulgaris genome assembly, annotation, proteome construction, and downstream analyses. Description of data: Manuscript-ready schematic summarizing the integrated bioinformatic workflow used for the S. vulgaris genomic and transcriptomic analyses. Panel A shows the primary workflow beginning with PacBio HiFi reads from a single migratory fifth-stage female larva, followed by adaptor screening, host depletion, contaminant screening, de novo assembly, repeat discovery and masking, transcript-supported annotation, proteome construction, and generation of discovery, representative, and comparative proteome datasets. Panel B shows the downstream analysis paths associated with each proteome resource, including functional annotation, secretome prediction, structure-guided annotation, transcriptomic and isoform analyses, orthology and gene-family evolution, putative HGT screening, resistance-homolog assessment, and the integrated outputs generated from these analyses.

**Additional file 2**

File format: Microsoft Word document (.docx)

Title of data: Additional file 2. Side-by-side comparison of the new Strongylus vulgaris genome assembly and the NCBI draft assembly.

Description of data: Comparative assembly statistics for the newly generated S. vulgaris genome assembly and the published NCBI draft assembly, including assembly size, scaffold and contig counts, contig N50, longest and mean contig lengths, GC content, BUSCO completeness categories, and estimated read depth.

**Additional file 3**

File format: Microsoft Word document (.docx)

Title of data: Additional file 3. Annotation and proteome refinement summary for Strongylus vulgaris.

Description of data: Summary of structural annotation and proteome refinement statistics for the S. vulgaris genome, including consensus transcript and locus counts, multi-transcript loci, novel loci, novel exons and introns, BUSCO annotation completeness, discovery proteome size, representative proteome size, isoforms removed, TE/repeat-associated proteins removed, and final comparative proteome size.

**Additional file 4**

File format: Microsoft Excel workbook (.xlsx)

Title of data: Additional file 4. Candidate secreted proteins of Strongylus vulgaris.

Description of data: Candidate secreted protein table for S. vulgaris, including protein identifiers, gene/locus and transcript identifiers, secretion-status classifications, signal-peptide and transmembrane-domain prediction fields, functional annotation status, and supporting metadata.

**Additional file 5**

File format: Microsoft Excel workbook (.xlsx)

Title of data: Additional file 5. Differential expression summary and top genes by contrast.

Description of data: Differential expression summary for pairwise life-stage and sex comparisons in S. vulgaris, including total genes tested, significant differentially expressed genes, direction of regulation, top upregulated genes, functional annotations, KO and Pfam support, baseMean values, log2 fold changes, adjusted P values, and sample-group metadata.

**Additional file 6**

File format: Microsoft Excel workbook (.xlsx)

Title of data: Additional file 6. Functional enrichment results by contrast.

Description of data: Functional enrichment results for S. vulgaris differential-expression contrasts, including KEGG pathway enrichment summaries, GSEA results, pathway identifiers, manuscript-safe pathway labels, enrichment statistics, driver genes, functional annotation support for driver genes, pathway-review labels, and source-file documentation.

**Additional file 7**

File format: Microsoft Excel workbook (.xlsx)

Title of data: Additional file 7. Isoform-switching and alternative-splicing summary.

Description of data: Isoform-switching and alternative-splicing results across S. vulgaris developmental and sex comparisons, including gene-level switch summaries, significant isoform-switch shortlist, isoform-level details, alternative-splicing event annotations, Iso-Seq support, isoform abundance values, transcript identifiers, isoform fractions, differential isoform usage statistics, and supporting metadata.

**Additional file 8**

File format: Microsoft Word document (.docx)

Title of data: Additional file 8. Comparative species and orthogroup summary.

Description of data: Comparative orthogroup summary for the eight-species nematode analysis, including species names, proteome sizes used in orthogroup analysis, assigned orthogroups, universal single-copy orthologues, shared orthogroups, and species-specific orthogroups. The table includes the final comparative S. vulgaris proteome and summarizes the S. vulgaris-specific orthogroup set.

**Additional file 9**

File format: Microsoft Excel workbook (.xlsx)

Title of data: Additional file 9. Expanded and contracted gene families in Strongylus vulgaris.

Description of data: CAFE5-derived gene-family expansion and contraction results for the S. vulgaris branch, including orthogroup identifiers, expansion/contraction status, CAFE5 P values, branch probabilities, inferred copy-number changes, species-level gene-copy counts, TE/repeat artifact flags, top expanded and contracted families, and CAFE5 model parameters.

**Additional file 10**

File format: Microsoft Excel workbook (.xlsx)

Title of data: Additional file10. Structure-guided annotation of weakly annotated proteins.

Description of data: Structure-guided prioritization of weakly annotated S. vulgaris proteins, including protein and locus identifiers, NCBI locus tags, transcript identifiers, Swiss-Prot and nematode homology status, Pfam/domain support, secretome status, differential-expression support, Foldseek and DeepFRI annotations, DALI and COFACTOR support where available, AlphaFold2 confidence categories, priority-candidate status, and evidence summaries.

**Additional file 11**

File format: Microsoft Excel workbook (.xlsx)

Title of data: Additional file 11. HGT screening and strict candidate summary.

Description of data: Horizontal gene transfer screening results for S. vulgaris, including all initial Alien Index candidates, best metazoan and non-metazoan matches, taxonomy and lineage information, RNA-seq support, scaffold and gene-structure metadata where available, GC content, codon-usage PCA coordinates, strict filtering outcomes, and the retained strict HGT candidate set.

**Additional file 12**

File format: Microsoft Excel workbook (.xlsx)

Title of data: Additional file 12. Resistance-associated homologs identified in Strongylus vulgaris.

Description of data: Summary of resistance-associated query proteins and their putative S. vulgaris homologs, including query species, drug class or resistance context, homolog identifiers, NCBI locus tags, percent identity, E values, orthogroup assignments, genomic-coordinate availability, copy-number interpretation, β-tubulin marker-site status, expression and Iso-Seq support, and phylogenetic tree availability for represented orthogroups.

## References

1. Kaplan RM, Nielsen MK. An evidence-based approach to equine parasite control: it ain’t the 60s anymore. Equine Vet Educ. 2010;22:306–316. doi:10.1111/j.2042-3292.2010.00084.x.

2. Reinemeyer CR, Nielsen MK. Handbook of equine parasite control. 2nd ed. Hoboken: John Wiley & Sons; 2018.

3. Bellaw JL, Nielsen MK. Meta-analysis of cyathostomin species-specific prevalence and relative abundance in domestic horses from 1975–2020: emphasis on geographical region and specimen collection method. Parasit Vectors. 2020;15. doi:10.1186/s13071-020-04396-5.

4. Love S, Duncan JL. Development of cyathostome infection of helminth-naive foals. Equine Vet J. 1992;24:93–98. doi:10.1111/j.2042-3306.1992.tb04796.x.

5. Wright AL. Verminous arteritis as a cause of colic in the horse. Equine Vet J. 1972;4:169–174. doi:10.1111/j.2042-3306.1972.tb03904.x.

6. Duncan JL. The life cycle, pathogenesis and epidemiology of S. vulgaris in the horse. Equine Vet J. 1973;5:20–25. doi:10.1111/j.2042-3306.1973.tb03188.x.

7. Hubert JD, Seahorn TL, Klei TR, Hosgood G, Horohov DW, Moore RM. Clinical signs and hematologic, cytokine, and plasma nitric oxide alterations in response to Strongylus vulgaris infection in helminth-naive ponies. Can J Vet Res. 2004;68:193–200.

8. Nielsen MK, Scare J, Gravatte HS, Bellaw JL, Prado JC, Reinemeyer CR. Changes in serum Strongylus vulgaris-specific antibody concentrations in response to anthelmintic treatment of experimentally infected foals. Front Vet Sci. 2015;2:17. doi:10.3389/fvets.2015.00017.

9. Pihl TH, Nielsen MK, Olsen SN, Leifsson PS, Jacobsen S. Nonstrangulating intestinal infarctions associated with Strongylus vulgaris: clinical presentation and treatment outcomes of 30 horses, 2008–2016. Equine Vet J. 2018;50:474–480. doi:10.1111/evj.12779.

10. White NA, Moore JN, Douglas M. SEM study of Strongylus vulgaris larva-induced arteritis in the pony. Equine Vet J. 1983;15:349–353. doi:10.1111/j.2042-3306.1983.tb01822.x.

11. Nielsen MK. Strongylus vulgaris – an ancient threat revisited. Equine Vet Educ. 2025;00:1–17. doi:10.1111/eve.70023.

12. Nielsen MK, Vidyashankar AN, Olsen SN, Monrad J, Thamsborg SM. Strongylus vulgaris associated with usage of selective therapy on Danish horse farms—Is it reemerging? Vet Parasitol. 2012. doi:10.1016/j.vetpar.2012.04.039.

13. Campbell BE, Nagaraj SH, Hu M, Zhong W, Sternberg PW, Ong EK, et al. Gender-enriched transcripts in Haemonchus contortus: predicted functions and genetic interactions based on comparative analyses with Caenorhabditis elegans. Int J Parasitol. 2008;38:65–83. doi:10.1016/j.ijpara.2007.07.001.

14. Cwiklinski K, Merga JY, Lake SL, Hartley C, Matthews JB, Paterson S, et al. Transcriptome analysis of a parasitic clade V nematode: comparative analysis of potential molecular anthelmintic targets in Cylicostephanus goldi. Int J Parasitol. 2013;43:917–927. doi:10.1016/j.ijpara.2013.06.010.

15. Lu MR, Lai CK, Liao BY, Tsai IJ. Comparative transcriptomics across nematode life cycles reveal gene expression conservation and correlated evolution in adjacent developmental stages. Genome Biol Evol. 2020;12:1019–1030. doi:10.1093/gbe/evaa110.

16. Coghlan A, Tyagi R, Cotton JA, Holroyd N, Rosa BA, Tsai IJ, et al. Comparative genomics of the major parasitic worms. Nat Genet. 2019;51:163–174. doi:10.1038/s41588-018-0262-1.

17. Coghlan A, Haegeman A. Comparative genomics of parasitic nematodes. Trends Genet. 2015;31:391–401.

18. Laing R, Kikuchi T, Martinelli A, Tsai IJ, Beech RN, Redman E, et al. Genome and transcriptome of Haemonchus contortus, a key model parasite for drug and vaccine discovery. Genome Biol. 2013;14:R88. doi:10.1186/gb-2013-14-8-r88.

19. Schwarz EM, Korhonen PK, Campbell BE, Young ND, Jex AR, Jabbar A, et al. The genome and developmental transcriptome of the strongylid nematode Haemonchus contortus. Genome Biol. 2013;14:R89. doi:10.1186/gb-2013-14-8-r89.

20. Doyle SR, Tracey A, Laing R, Holroyd N, Bartley D, Bazant W, et al. Genomic and transcriptomic variation defines the chromosome-scale assembly of Haemonchus contortus, a model gastrointestinal worm. Commun Biol. 2020;3:656. doi:10.1038/s42003-020-01377-3.

21. Pertea M, Pertea GM, Antonescu CM, Chang TC, Mendell JT, Salzberg SL. StringTie enables improved reconstruction of a transcriptome from RNA-seq reads. Nat Biotechnol. 2015;33:290–295. doi:10.1038/nbt.3122.

22. Marx V. Method of the year: long-read sequencing. Nat Methods. 2023;20:6–11. doi:10.1038/s41592-022-01730-w.

23. Barberan-Soler S, Zahler AM. Alternative splicing regulation during C. elegans development: splicing factors as regulated targets. PLoS Genet. 2008;4:e1000001. doi:10.1371/journal.pgen.1000001.

24. Barberan-Soler S, Lambert NJ, Zahler AM. Global analysis of alternative splicing uncovers developmental regulation of nonsense-mediated decay in C. elegans. RNA. 2009;15:1652–1660. doi:10.1261/rna.1711109.

25. Britton C, Roberts B, Marks ND. Functional genomics tools for Haemonchus contortus and lessons from other helminths. Adv Parasitol. 2016;93:599–623. doi:10.1016/bs.apar.2016.02.017.

26. Jones JT, Smant G, Blok VC. SXP/RAL-2 proteins of the potato cyst nematode Globodera rostochiensis: secreted proteins of the hypodermis and amphids. Nematology. 2000;2:887–893. doi:10.1163/156854100750112833.

27. Tytgat T, Vercauteren I, Vanholme B, De Meutter J, Vanhoutte I, Gheysen G, et al. An SXP/RAL-2 protein produced by the subventral pharyngeal glands in the plant parasitic root-knot nematode Meloidogyne incognita. Parasitol Res. 2005;95:50–54. doi:10.1007/s00436-004-1243-0.

28. Lozano-Torres JL, Wilbers RHP, Warmerdam S, Finkers-Tomczak A, Diaz-Granados A, van Schaik CC, et al. Apoplastic venom allergen-like proteins of cyst nematodes modulate the activation of basal plant innate immunity by cell surface receptors. PLoS Pathog. 2014;10:e1004569. doi:10.1371/journal.ppat.1004569.

29. Wilbers RHP, Schneiter R, Holterman MHM, Drurey C, Smant G, Asojo OA, et al. Secreted venom allergen-like proteins of helminths: conserved modulators of host responses in animals and plants. PLoS Pathog. 2018;14:e1007300. doi:10.1371/journal.ppat.1007300.

30. Maizels RM, Smits HH, McSorley HJ. Modulation of host immunity by helminths: the expanding repertoire of parasite effector molecules. Immunity. 2018;49:801–818. doi:10.1016/j.immuni.2018.10.016.

31. Bouchery T, Moyat M, Sotillo J, Silverstein S, Volpe B, Coakley G, et al. Hookworms evade host immunity by secreting a deoxyribonuclease to degrade neutrophil extracellular traps. Cell Host Microbe. 2020;27:277–289.e6. doi:10.1016/j.chom.2020.01.011.

32. Haegeman A, Jones JT, Danchin EGJ. Horizontal gene transfer in nematodes: a catalyst for plant parasitism? Mol Plant Microbe Interact. 2011;24:879–887. doi:10.1094/MPMI-03-11-0055.

33. Danchin EGJ, Guzeeva EA, Mantelin S, Berepiki A, Jones JT. Horizontal gene transfer from bacteria has enabled the plant-parasitic nematode Globodera pallida to feed on host-derived sucrose. Mol Biol Evol. 2016;33:1571–1579. doi:10.1093/molbev/msw041.

34. Li J, Xu C, Yang S, Chen C, Tang S, Wang J, et al. A venom allergen-like protein, RsVAP, the first discovered effector protein of Radopholus similis that inhibits plant defense and facilitates parasitism. Int J Mol Sci. 2021;22:4782. doi:10.3390/ijms22094782.

35. Chang Q, Yang Y, Hong B, Zhao Y, Zhao M, Han S, et al. A variant of the venom allergen-like protein, DdVAP2, is required for the migratory endoparasitic plant nematode Ditylenchus destructor parasitism of plants. Front Plant Sci. 2023;14:1322902. doi:10.3389/fpls.2023.1322902.

36. Gong H, Kobayashi K, Sugi T, Takemae H, Kurokawa H, Horimoto T, et al. A novel PAN/apple domain-containing protein from Toxoplasma gondii: characterization and receptor identification. PLoS One. 2012;7:e30169. doi:10.1371/journal.pone.0030169.

37. Paoletta MS, Wilkowsky SE. Thrombospondin related anonymous protein superfamily in vector-borne apicomplexans: the parasite’s toolkit for cell invasion. Front Cell Infect Microbiol. 2022;12:831592. doi:10.3389/fcimb.2022.831592.

38. Jumper J, Evans R, Pritzel A, Green T, Figurnov M, Ronneberger O, et al. Highly accurate protein structure prediction with AlphaFold. Nature. 2021;596:583–589. doi:10.1038/s41586-021-03819-2.

39. Gligorijević V, Renfrew PD, Kosciolek T, Koehler Leman J, Berenberg D, Vatanen T, et al. Structure-based protein function prediction using graph convolutional networks. Nat Commun. 2021;12:3168. doi:10.1038/s41467-021-23303-9.

40. van Kempen M, Kim SS, Tumescheit C, Mirdita M, Lee J, Gilchrist CLM, et al. Fast and accurate protein structure search with Foldseek. Nat Biotechnol. 2024;42:243–246. doi:10.1038/s41587-023-01773-0.

41. Paganini J, Campan-Fournier A, Da Rocha M, Gouret P, Pontarotti P, Wajnberg E, et al. Contribution of lateral gene transfers to the genome composition and parasitic ability of root-knot nematodes. PLoS One. 2012;7:e50875. doi:10.1371/journal.pone.0050875.

42. Nielsen MK. Anthelmintic resistance in equine nematodes: current status and emerging trends. Int J Parasitol Drugs Drug Resist. 2022;20:76–88. doi:10.1016/j.ijpddr.2022.10.005.

43. Hodgkinson JE, Clark HJ, Kaplan RM, Lake SL, Matthews JB. The role of polymorphisms at β-tubulin isotype 1 codons 167 and 200 in benzimidazole resistance in cyathostomins. Int J Parasitol. 2008;38:1149–1160. doi:10.1016/j.ijpara.2008.02.001.

44. Khan S, Nisar A, Yuan J, Luo X, Dou X, Liu F, et al. A whole genome re-sequencing based GWA analysis reveals candidate genes associated with ivermectin resistance in Haemonchus contortus. Genes. 2020;11:367. doi:10.3390/genes11040367.

45. Doyle SR, Laing R, Bartley D, Morrison A, Holroyd N, Maitland K, et al. Genomic landscape of drug response reveals mediators of anthelmintic resistance. Cell Rep. 2022;41:111522. doi:10.1016/j.celrep.2022.111522.

46. American Veterinary Medical Association. AVMA guidelines for the euthanasia of animals: 2020 edition. Schaumburg: American Veterinary Medical Association; 2020.

47. Lichtenfels JR, Kharchenko VA, Dvojnos GM. Illustrated identification keys to strongylid parasites (Strongylidae: Nematoda) of horses, zebras and asses (Equidae). Vet Parasitol. 2008;156:4–161. doi:10.1016/j.vetpar.2008.04.026.

48. Sim SB, Corpuz RL, Simmonds TJ, Geib SM. HiFiAdapterFilt, a memory efficient read processing pipeline, prevents occurrence of adapter sequence in PacBio HiFi reads and their negative impacts on genome assembly. BMC Genomics. 2022;23(1):157. doi:10.1186/s12864-022-08375-1.

49. National Center for Biotechnology Information. Genome assembly TB-T2T [Internet]. Bethesda: National Center for Biotechnology Information; 2024 [Accessed: 2025 March 22]. Available from: https://www.ncbi.nlm.nih.gov/datasets/genome/GCF_041296265.1/

50. Rhie A, Walenz BP, Koren S, Phillippy AM. Merqury: reference-free quality, completeness, and phasing assessment for genome assemblies. Genome Biol. 2020;21:245. doi:10.1186/s13059-020-02134-9.

51. Ranallo-Benavidez TR, Jaron KS, Schatz MC. GenomeScope 2.0 and Smudgeplot for reference-free profiling of polyploid genomes. Nat Commun. 2020;11(1):1432. doi:10.1038/s41467-020-14998-3.

52. Cheng H, Concepcion GT, Feng X, Zhang H, Li H. Haplotype-resolved de novo assembly using phased assembly graphs with hifiasm. Nat Methods. 2021;18:170–175. doi:10.1038/s41592-020-01056-5.

53. Manni M, Berkeley MR, Seppey M, Simão FA, Zdobnov EM. BUSCO update: novel and streamlined workflows along with broader and deeper phylogenetic coverage for scoring of eukaryotic, prokaryotic, and viral genomes. Mol Biol Evol. 2021;38:4647–4654. doi:10.1093/molbev/msab199.

54. Smit AFA, Hubley R, Green P. RepeatMasker Open-4.0 [software]. 2013.

55. Bruna T, Hoff KJ, Lomsadze A, Stanke M, Borodovsky M. BRAKER3: fully automated genome annotation using RNA-seq and protein evidence with GeneMark-ETP, AUGUSTUS, and TSEBRA. Genome Res. 2024;34:855–867. doi:10.1101/gr.278373.123.

56. Emms DM, Kelly S. OrthoFinder: phylogenetic orthology inference for comparative genomics. Genome Biol. 2019;20:238. doi:10.1186/s13059-019-1832-y.

57. Mendes FK, Vanderpool D, Fulton B, Hahn MW. CAFE 5 models variation in evolutionary rates among gene families. Bioinformatics. 2020;36:5516–5518. doi:10.1093/bioinformatics/btaa1022.

58. Moriya Y, Itoh M, Okuda S, Yoshizawa AC, Kanehisa M. KAAS: an automatic genome annotation and pathway reconstruction server. Nucleic Acids Res. 2007;35:W182–W185. doi:10.1093/nar/gkm321.

59. Teufel F, Almagro Armenteros JJ, Johansen AR, Gíslason MH, Pihl SI, Tsirigos KD, et al. SignalP 6.0 predicts all five types of signal peptides using protein language models. Nat Biotechnol. 2022;40:1023–1025. doi:10.1038/s41587-021-01156-3.

60. Kall L, Krogh A, Sonnhammer ELL. A combined transmembrane topology and signal peptide prediction method. J Mol Biol. 2004;338:1027–1036. doi:10.1016/j.jmb.2004.03.016.

61. Benjamini Y, Hochberg Y. Controlling the false discovery rate: a practical and powerful approach to multiple testing. J R Stat Soc Series B Stat Methodol. 1995;57:289–300. doi:10.1111/j.2517-6161.1995.tb02031.x.

62. Wu T, Hu E, Xu S, Chen M, Guo P, Dai Z, et al. clusterProfiler 4.0: a universal enrichment tool for interpreting omics data. Innovation. 2021;2:100141. doi:10.1016/j.xinn.2021.100141.

63. Vitting-Seerup K, Sandelin A. IsoformSwitchAnalyzeR: analysis of changes in genome-wide patterns of alternative splicing and its functional consequences. Bioinformatics. 2019;35:4469–4471. doi:10.1093/bioinformatics/btz247.

64. Minh BQ, Schmidt HA, Chernomor O, Schrempf D, Woodhams MD, von Haeseler A, et al. IQ-TREE 2: new models and efficient methods for phylogenetic inference in the genomic era. Mol Biol Evol. 2020;37:1530–1534. doi:10.1093/molbev/msaa015.

65. Hawdon JM, Jones BF, Hoffman DR, Hotez PJ. Cloning and characterization of Ancylostoma-secreted protein: a novel protein associated with the transition to parasitism by infective hookworm larvae. J Biol Chem. 1996;271:6672–6678. doi:10.1074/jbc.271.12.6672.

66. Mohandas N, Young ND, Jabbar A, Korhonen PK, Koehler AV, Amani P, et al. The barber’s pole worm CAP protein superfamily: a basis for fundamental discovery and biotechnology advances. Biotechnol Adv. 2015;33:1744–1754. doi:10.1016/j.biotechadv.2015.07.003.

67. Doyle SR, Illingworth CJR, Laing R, Bartley DJ, Redman E, Martinelli A, et al. Population genomic and evolutionary modelling analyses reveal a single major QTL for ivermectin drug resistance in the pathogenic nematode Haemonchus contortus. BMC Genomics. 2019;20:218. doi:10.1186/s12864-019-5592-6.

