## Additional File 2 for "Comparative genomic and transcriptomic analyses of Strongylus vulgaris reveal developmental and evolutionary deployment of parasitism in a migratory equine nematode"

**Additional File 1.** Side-by-side comparison of the new *Strongylus vulgaris* genome assembly and the NCBI draft assembly.

| **Metric** | **New assembly** | **NCBI draft assembly** |
| --- | --- | --- |
| Assembly size | 329,228,623 bp (329.2 Mb) | 291.1 Mb |
| Number of scaffolds | 3,913 | 167,310 |
| Number of contigs | 3,913 | 397,821 |
| Contig N50 | 154 kb | 939 bp |
| Longest contig | 961 kb | 88 kb |
| Mean contig length | 84 kb | 732 bp |
| GC content | 38.66% | 38.66% |
| BUSCO complete (C) | 94.2% (2,950/3,131) | 21.7% |
| BUSCO single-copy (S) | 84.9% (2,657/3,131) | 11.4% |
| BUSCO duplicated (D) | 9.4% (293/3,131) | 10.3% |
| BUSCO fragmented (F) | 3.5% (109/3,131) | 15.2% |
| BUSCO missing (M) | 2.3% (72/3,131) | 63.1% |
| Estimated read depth | 105x | 43x |

*Note.* Mean contig length was calculated as assembly size divided by number of contigs. BUSCO values are reported from nematoda_odb10 (n = 3,131). *Abbreviations:* BUSCO, Benchmarking Universal Single-Copy Orthologs.
