## Additional File 3 for "Comparative genomic and transcriptomic analyses of Strongylus vulgaris reveal developmental and evolutionary deployment of parasitism in a migratory equine nematode"

**Additional file 2.** Annotation and proteome refinement summary for *Strongylus vulgaris*. Structural annotation statistics were derived from the final transcript GTF comparison, and proteome counts summarize isoform collapsing and repeat/TE filtering used to generate the final comparative proteome.

| **Analysis component** | **Metric** | **Value** |
| --- | --- | --- |
| Structural annotation | Transcript count | 24,546 consensus transcripts |
| Structural annotation | Locus count | 16,387 loci |
| Structural annotation | Multi-transcript loci | 4,607 loci (~1.5 transcripts per locus) |
| Structural annotation | Novel loci | 1,259 of 16,387 loci (7.7%) |
| Structural annotation | Novel exons | 15,556 of 168,096 exons (9.3%) |
| Structural annotation | Novel introns | 13,557 of 148,092 introns (9.2%) |
| Annotation completeness | BUSCO annotation completeness | C: 90.9% [S: 50.3%, D: 40.5%], F: 4.5%, M: 4.6%, n = 16,387 |
| Proteome refinement | Discovery proteome size | 23,363 proteins |
| Proteome refinement | Representative proteome size | 17,171 proteins |
| Proteome refinement | Isoforms removed | 6,192 isoform entries |
| Proteome refinement | TE/repeat-associated proteins removed | 786 proteins |
| Proteome refinement | Final comparative proteome size | 16,385 proteins |

**Note.** Discovery proteome size refers to the full TransDecoder-derived protein set prior to isoform collapsing. The representative proteome was collapsed to one protein per locus before removal of TE/repeat-associated proteins to generate the final comparative proteome.
