## Additional File 8 for "Comparative genomic and transcriptomic analyses of Strongylus vulgaris reveal developmental and evolutionary deployment of parasitism in a migratory equine nematode"

**Additional file 7.** Comparative species and orthogroup summary.

| **Species name** | **Source/reference** | **Proteome size used†** | **Assigned orthogroups** | **Single-copy orthologues‡** | **Shared orthogroups§** | **Species-specific orthogroups¶** |
| --- | --- | --- | --- | --- | --- | --- |
| *Caenorhabditis elegans* | Public proteome; source metadata pending | 25,614 | 11,333 | 91 | 9,430 | 1,903 |
| *Nippostrongylus brasiliensis* | Public proteome; source metadata pending | 27,889 | 12,894 | 91 | 11,895 | 999 |
| *Haemonchus contortus* | Public proteome; source metadata pending | 32,784 | 12,516 | 91 | 11,405 | 1,111 |
| *Oesophagostomum dentatum* | Public proteome; source metadata pending | 22,299 | 12,143 | 91 | 11,315 | 828 |
| *Cylicocyclus nassatus* | Public proteome; source metadata pending | 20,247 | 12,471 | 91 | 11,640 | 831 |
| *Necator americanus* | Public proteome; source metadata pending | 37,900 | 14,317 | 91 | 12,512 | 1,805 |
| *Ancylostoma ceylanicum* | Public proteome; source metadata pending | 59,081 | 18,478 | 91 | 14,061 | 4,417 |
| *Strongylus vulgaris* | This study; final comparative proteome | 15,348 | 10,348 | 91 | 10,147 | 201 |

**Notes.** † Values are proteins represented in Orthogroups.tsv, meaning proteins assigned to orthogroups. True total input proteome sizes and unassigned proteins should be updated when the regenerated OrthoFinder per-species statistics or original FASTA summaries are available.

‡ Single-copy orthologues are defined here as universal one-to-one orthogroups containing exactly one protein from each of the eight species.

§ Shared orthogroups are orthogroups present in the focal species and at least one additional species.

¶ Species-specific orthogroups are orthogroups present only in the focal species within this eight-species comparison. The *Strongylus vulgaris*-specific set comprises 201 orthogroups and 640 proteins.
