## Supplementary figures and images for "Comparative genomic and transcriptomic analyses of Strongylus vulgaris reveal developmental and evolutionary deployment of parasitism in a migratory equine nematode"

### Additional File 1

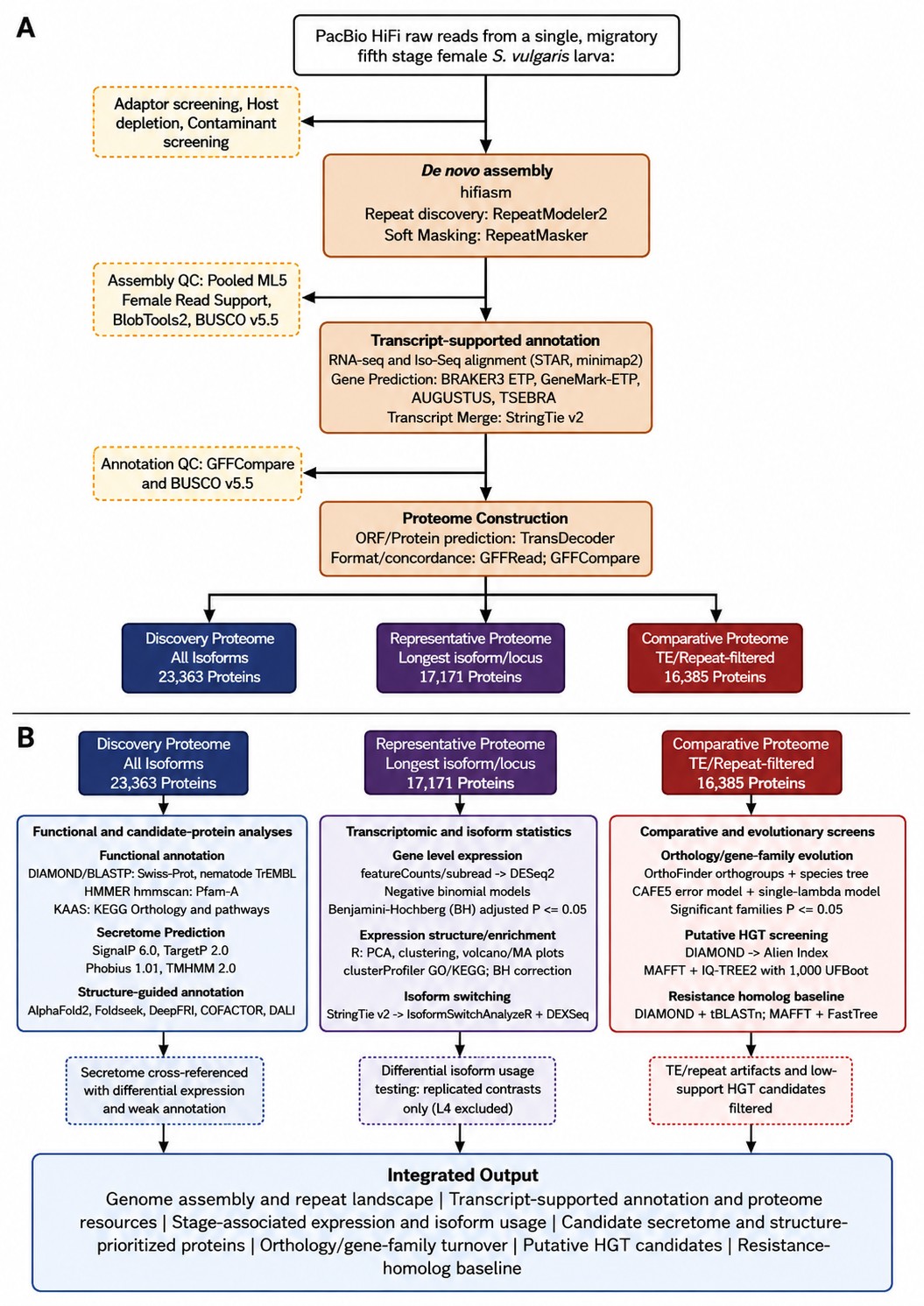
